# Requirements for development of T-helper1 and T-follicular helper cells from a common precursor

**DOI:** 10.1101/2025.09.08.674857

**Authors:** Douwe M. T. Bosma, Julia Busselaar, Mo D. Staal, Mylene de Koning, Xin Lei, Tom de Wit, Yanling Xiao, Jannie Borst, Fiamma Salerno

## Abstract

Activated CD4 T cells become either T-helper (Th) or T-follicular helper (Tfh) cells that support cellular or humoral immunity. We have investigated how polyclonal, vaccine induced T cells in mice bifurcate into Th1- and Tfh trajectories. We show that Th1 and Tfh cells originate from the same, highly proliferative precursor clones that co-express Th1- and Tfh-related transcription factors and chemokine receptors, including T-bet, BCL6, CXCR3 and CXCR5. Generation of the common Th1/Tfh precursor pool from antigen-specific CD4 T cells relies on CD28 costimulation but occurs independently of type-1 conventional dendritic cells (cDC1s) or B cells. Differentiation of Th1/Tfh precursors into the Th1 lineage relies on CD40 costimulation and cDC1s, while differentiation into the Tfh lineage relies on ICOS costimulation and B cells. Thus, activated CD4 T cells give rise to a bipotent Th1/Tfh precursor and differentiation into Th1 or Tfh cells depends on interactions with antigen-presenting cDC1s or B cells.

## Introduction

Upon antigen encounter within secondary lymphoid organs, CD4 T cells are activated, clonally expand and initiate differentiation into T-helper (Th)1, 2 or 17 lineages to regulate cellular immunity (Yin et al., 2021). At the same time, T-follicular helper (Tfh) cells are formed, which promote B-cell expansion and differentiation in germinal centers (GC) (Vinuesa et al., 2016). During a type-I immune response, Th1 and Tfh cells emerge in parallel, suggesting a bifurcation in their effector differentiation trajectories (Duckworth & Groom, 2021; Lönnberg et al., 2017; Osum & Jenkins, 2023). This process is not exclusive of type-I responses, as Tfh cells are likewise generated in parallel with Th2 or Th17 effector CD4 T cells (Eisenbarth et al., 2021; Künzli & Masopust, 2023). Coordination of cell-mediated and humoral immunity thus depends on Th/Tfh lineage decision making in activated CD4 T cells. This process is a potential target for therapeutic intervention e.g. in vaccination, cancer and auto-immunity.

Effector CD4 T-cell differentiation relies on successive interactions with different professional antigen-presenting cells (APCs) and is initiated upon cognate contact with conventional dendritic cells (cDCs). Among these, the cDC2 lineage excels at (soluble) antigen presentation in major histocompatibility complex (MHC)-II molecules (Dudziak et al., 2007) and are important for initial CD4 T-cell activation (De Giovanni et al., 2020; Hilligan & Ronchese, 2020). The cDC1 lineage is much more proficient in phagocytosing cell-bound antigen and cross-presenting this in MHC-I molecules, thus playing a crucial role in CD8 T-cell activation (Den Haan et al., 2000; Hildner et al., 2008; Martínez-López et al., 2015). cDC1s can also contribute to CD4 T cell priming (Ferris et al., 2020; Wu & Murphy, 2022) and relay help for effector and memory differentiation of CD8 T cells (Borst et al., 2018; Lei et al., 2024). In contrast, B cells are considered essential APCs to promote CD4 T-cell differentiation into Tfh cells (Crotty, 2014; Lönnberg et al., 2017).

In general, effector differentiation of CD4 T cells is guided by the activation and maturation state of cDCs and B cells, which are reflected in their costimulatory and cytokine profiles (De Giovanni et al., 2020; Künzli & Masopust, 2023; Yin et al., 2021). These inputs, as delivered during multiple sequential interactions between T cells and APCs collectively regulate expression of lineage-determining transcription factors that direct Th differentiation along specific trajectories. Decision-making between Th and Tfh lineage commitment likewise depends on certain lineage-defining transcription factors. Whereas transient co-expression of T-bet and BCL6 has been reported in *in vitro* T-cell cultures (Nakayamada et al., 2011), *in vivo* studies indicate that the choice between Th1 and Tfh cell differentiation depends on the ratio of T-bet versus BCL6 protein levels, which is determined by initiation of negative regulatory circuits. For example, BCL6 represses genes encoding Th1, Th2 and Th17 lineage-specific transcription factors, including *Tbx21* (encoding T-bet), *Gata3* and *Rora* (J. Choi et al., 2020). However, when it binds to T-bet, BCL6 can no longer repress the *Prdm1* gene encoding BLIMP1. As a result, BLIMP1 starts to suppress Tfh-associated genes, thus favoring Th1 differentiation (Oestreich et al., 2012). In contract, transient early T-bet expression is required for upregulation of CXCR5 and generation of IFNγ-producing Tfh cells in response to infection (Fang et al., 2018; Weinstein et al., 2018).

In this paper, we address the mechanisms underlying bifurcation of the Th1 and Tfh lineages. The CD4 T-cell differentiation trajectories leading to these cell fates have generally been studied separately (Osum & Jenkins, 2023). Also, they have primarily focused on T cell receptor (TCR)-transgenic CD4 T cells (Cho et al., 2017; Fazilleau et al., 2009; Lönnberg et al., 2017; Tubo et al., 2013) and not yet revealed the bifurcation process during endogenous, polyclonal CD4 T-cell responses (Künzli & Masopust, 2023). This is a relevant distinction since TCR affinity has been proposed to impact the differentiation trajectories in some studies (Fazilleau et al., 2009; Künzli et al., 2021; Tubo et al., 2013), but not in others (Cho et al., 2017; De Giovanni et al., 2020). Deconvolution of CD4 T-cell differentiation pathways can be challenging due to high plasticity of CD4 T-helper lineages (Alterauge et al., 2020; O’Shea & Paul, 2010) and stochastic events (Kiner et al., 2021).

Different lines of evidence point to the existence of a common precursor of Th1 and Tfh cells. Single cell RNA-sequencing (scRNAseq) of antigen-specific TCR transgenic PbTII CD4 T cells to *Plasmodium chabaudi* revealed that prior to Th1 or Tfh differentiation, CD4 T cells attain a common activated precursor state co-expressing *Cxcr5* and *Cxcr3* mRNA (Lönnberg et al., 2017). In lymphocytic choriomeningitis virus (LCMV), a memory-like CD4 precursor was found to be clonally related to Th1 and Tfh cells (Xia et al., 2022). Understanding whether and how overlapping Th1/Tfh intermediate cell states arise in normal polyclonal responses is critical, as these cells represent a promising target to therapeutically direct Th1 or Tfh differentiation in disease. Defining the signals that bias differentiation toward either lineage is therefore essential. These precursors also hold potential as prognostic or predictive markers and for immunomonitoring of treatment responses.

Here, we used a mouse vaccination model to induce and trace the differentiation trajectories of endogenous CD4 T cells in healthy mice. By combining high-dimensional flow cytometry, single-cell RNA-sequencing (scRNAseq) and TCR-sequencing, we identified that antigen-specific CD4 T clones pass through a bipotent Th1/Tfh precursor state, before committing to a specific differentiation trajectory. We identify the gene and protein expression profile of this precursor and extrapolate its existence to mouse infection settings. We also define key costimulatory signals and interactions with APCs that guide differentiation commitment of the Th1/Tfh precursor to either the Th1 or Tfh lineage.

## Results

### Vaccination raises endogenous, functional Th1 and Tfh cells in equal measure

To study how endogenous CD4 T cells differentiate *in vivo*, we used a mouse vaccination model that has previously revealed the effects of CD4 T-cell help on effector and memory CD8 T-cell differentiation (Ahrends et al., 2016, 2017, 2019). In this model, we vaccinate C57BL/6 mice with plasmid DNA encoding an immunodominant MHC-I (H-2D^b^) restricted epitope E7_48-57_ from human papilloma virus (HPV) either alone (Control) or linked to three MHC-II-restricted helper epitopes: p30, PADRE, and NEF (MHC-II) (**Figure 1A**). These MHC-II-restricted epitopes were selected for binding to human HLA-DP, -DQ and -DR, respectively (Oosterhuis et al., 2012), but p30 and PADRE can also be presented by MHC-II (I-A^b^) (Alexander et al., 1994; Rice et al., 2001) and prime CD4 T-cell responses in mice (Ahrends et al., 2016; Bosma et al., 2025). The plasmid DNA vaccine is “tattooed” into the depilated skin of the hind leg and expressed in keratinocytes, followed by antigen delivery to the draining lymph node (dLN) by cDCs and by passive drainage (Babała et al., 2018; Bins et al., 2005). The antigen-specific CD4 T-cell response can be followed by flow cytometry using an MHC-II tetramer (tet) against the PADRE epitope (**Figure 1B**). PADRE tet^+^ CD4 T cells were detected in the dLN from day 4 onwards after vaccination (**Figure 1C**) and entered the circulation by day 5 (**Supplementary** Figure 1A). In line with a type-1 immune response raised by this model, PADRE tet^+^ cells differentiated into Th1 and Tfh cells, and did not differentiate into T-regulatory (Treg), Th2 or Th17 cells (**Supplementary** Figure 1B). Among PADRE tet^+^ CD4 T cells, Th1 cells were defined by PSGL1, T-bet and SLAM coexpression and low or no TCF1 expression (**Figure 1D; Supplementary** Figure 1C) (Marshall et al., 2011; Shaw et al., 2016; Zou et al., 2024). Conversely, Tfh cells were defined by low or no PSGL1, T-bet and SLAM expression, gain of CXCR5, BCL6, PD-1 and maintained TCF1 expression (**Figure 1D**) (Y. S. Choi et al., 2015; Künzli & Masopust, 2023; Marshall et al., 2011; Osum & Jenkins, 2023). Th1 and Tfh populations emerged in parallel and were of similar size at day 4 after vaccination (**Figure 1E**). By day 5, when PADRE tet^+^ CD4 T cells became detectable in the blood (**Supplementary** Figure 1A), Tfh cells started accumulating in the dLN (**Figure 1E**). At that day, Th1 cells went from the dLN to the blood, where they represented ∼50% of PADRE^+^ CD4 T cells (**Figure 1F**).

**Figure 1:**
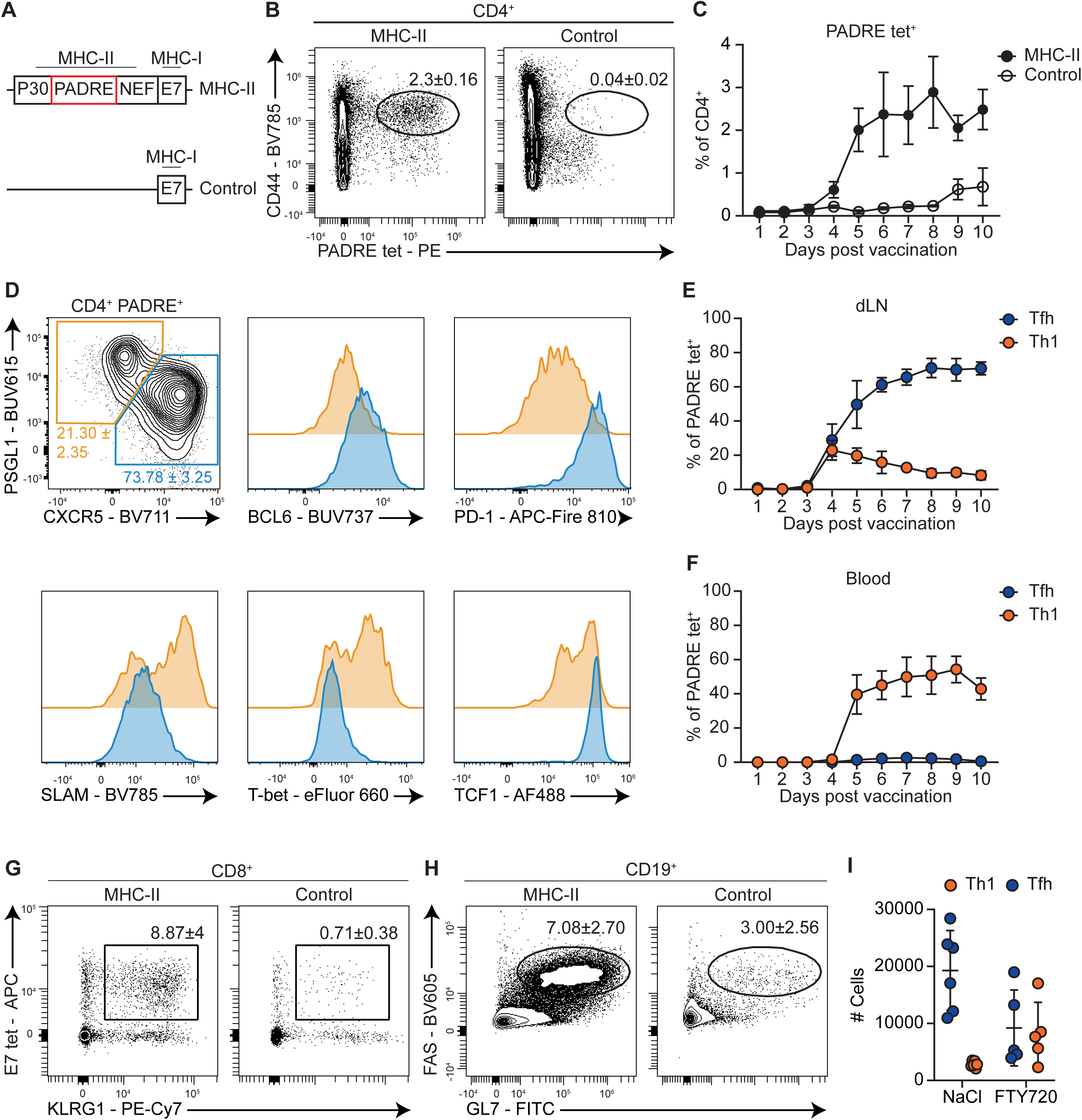
A vaccination model to study differentiation of vaccine-primed CD4 T cells into Th1 and Tfh cells. A) Scheme of epitopes in MHC-II and control DNA vaccines. B) Representative flow cytometry plots depicting frequency of CD44^+^ PADRE tet^+^ CD4 T cells in dLN at day 5 post-vaccination (N=3, mean ± SD). C) Frequency of PADRE tet^+^ CD4 T cells in dLN over time (N=8 for MHC-II, N=3 for Control). D) Representative flow cytometry plots depicting the phenotype of PADRE tet^+^ CD4 T cells in the dLN at day 8 post-vaccination. Colors in histograms match the CXCR5^-^ PSGL1^hi^ (orange) or CXCR5^+^ PSGL1^mid-lo^ (blue) populations E, F) Frequency of Tfh (CXCR5^+^ PSGL1^lo^ PD-1^+^ BCL6^+^) and Th1 cells (CXCR5^-^ PSGL1^hi^ T-bet^+^ SLAM^+^) within PADRE tet^+^ CD4 T cells in dLN (E) or blood (F) over time after MHC-II-vaccination (N=5). G) Representative contour plots depicting antigen-specific effector CD8 T cells (E7 tet^+^ KLRG1^+^) in blood at day 10 after MHC-II or control vaccination. H) Representative contour plots depicting GL7^+^ FAS^+^ GC B cells in the dLN at day 8 after MHC-II or control vaccination. I) Absolute numbers of Th1 and Tfh PADRE tet^+^ CD4 T cells in dLN at d7 post vaccination (N=6 mean ± SD).

To validate the helper functions of Th1 and Tfh cells, we investigated the CD8 T-cell and B-cell responses post vaccination. Mice in which CD4 T cells were activated by the MHC-II vaccine displayed an increased frequency of KLRG1^+^ effector phenotype E7 tetramer^+^ CD8 T cells at day 10 in the blood compared to mice receiving the control vaccine (**Figure 1G; Supplementary figure 1D**). These data confirm that the Th1 cells generated by the MHC-II vaccine contributed to effector differentiation of the CD8 T cells raised by the E7 epitope, as demonstrated before (Ahrends et al., 2016, 2017). Mice that received the MHC-II vaccine also displayed increased frequencies of FAS^+^ GL7^+^ germinal center (GC) B cells and CD138^+^ TACI^+^ plasmablasts (**Figure 1H; Supplementary** Figure 1E-F), confirming that vaccine-primed Tfh cells promoted the B-cell response. Thus, both Th1 and Tfh cells raised by the MHC-II vaccine are functionally competent. Inhibition of T-cell egress from the dLN with sphingosine 1-phosphate (S1P) receptor-targeting drug FTY720 increased the overall frequency of PADRE tet^+^ CD4 T cells in the dLN by more than 2-fold (**Supplementary** Figure 1G**, H**). At day 7 after vaccination, PADRE tet^+^ Th1 and Tfh cell frequencies and absolute numbers were comparable in dLN of FTY720-treated mice (**Figure 1I; Supplementary** Fig 1I). This finding indicates that the MHC-II vaccine leads in equal measure to Th1 and Tfh differentiation and therefore is a good model to study the differentiation trajectories of the endogenous vaccine-specific CD4 T cells raised.

### Reconstruction of the Th1- and Tfh differentiation trajectories that evolve after vaccination

To gain insight into the temporal changes that CD4 T cells undergo during Th1 and Tfh differentiation, we analyzed PADRE tet^+^ CD4 T cells in the dLNs of mice from day 4 to 9 after vaccination by spectral flow cytometry and dimension reduction analysis (**Figure 2A; Supplementary** Figure 2A). To ensure that we captured the full differentiation trajectory, we gated on total CD4^+^ PADRE tet^+^ cells, which include a small proportion of CD62L^hi^ CD44^low^ naïve cells. Indeed, PADRE tet^+^ CD4 T cells were discriminated into naïve CD62L^hi^ CD44^low^ and activated CD44^hi^ cells (**Figure 2B**). The marker profile of the activated PADRE^+^ CD4 T-cell pool underwent daily remodeling and displayed progressive branching towards two end points (**Figure 2A; Supplementary** Figure 2A), which were characterized by a Th1 (PSGL1^+^ T-bet^+^ SLAM^+^ TCF1^-^ SLAMF6^-^) or a Tfh (CXCR5^+^ PD-1^+^ BCL6^+^ TCF1^+^ SLAMF6^+^ FR4^+^) phenotype (**Figure 2C-E**), as we had observed before (**Figure 1D**). To reconstruct the temporal evolution of PADRE tet^+^ CD4 T-cell differentiation towards Th1 and Tfh states, we used Wishbone, an algorithm that orders cells according to their developmental progression and infers branching trajectories into only one of two possible fates (Setty et al., 2016). Wishbone calculates differentiation trajectories based on the rise and fall of phenotypic markers and uses nearest-neighbor graphs to capture developmental distance from a starting point, which we defined as the CD62L^hi^ CD44^low^ naive CD4 T-cell state.

**Figure 2:**
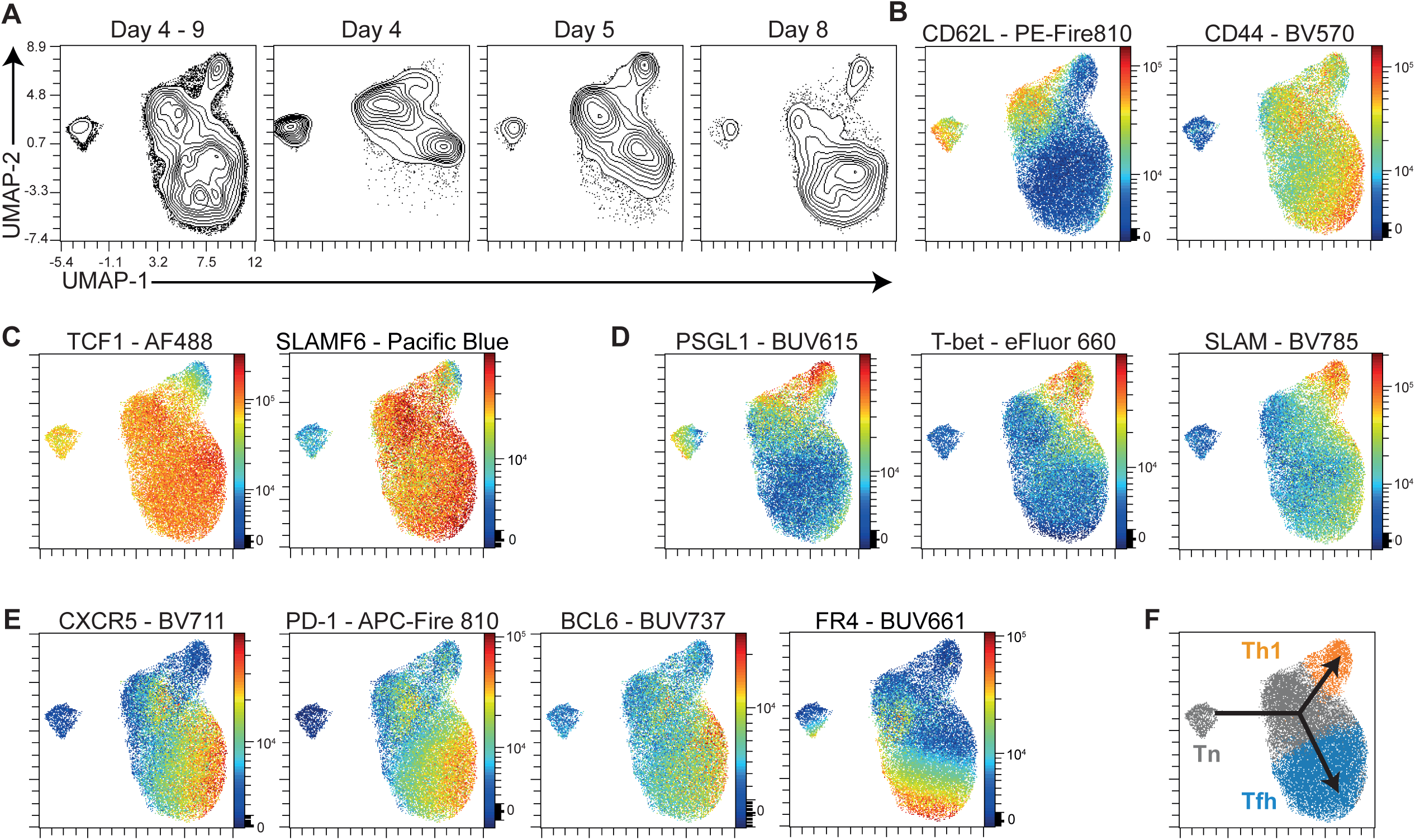
Temporal analysis highlights a common differentiation branchpoint for Th1 and Tfh cells. A) UMAP projections showing concatenated PADRE tet^+^ CD4 T cells in dLN from day 4-9 post-vaccination, or single projections at days 4, 5 or 8 (N=5 per time point). B-E) Overlay of CD62L and CD44 (B), TCF1 and SLAMF6 (C), PSGL1, T-bet and SLAM (D) and CXCR5, PD-1, BCL6 and FR4 expression (E). F) Overlay of Wishbone branching on UMAP from panel D. Arrows depict differentiation branches. Grey: trunk; orange: branch 1, identified as Th1 cells; blue: branch 2, identified as Tfh cells.

In our experimental setup, PADRE tet^+^ CD4 T cells were ordered based on early activation (CD44, CD62L) and stem-like markers (TCF1, SLAMF6) and the acquisition of the Th1 or Tfh phenotype (Y. S. Choi et al., 2015; Kageyama et al., 2012; Künzli et al., 2020; Marshall et al., 2011; Osum & Jenkins, 2023; Shaw et al., 2016; Zou et al., 2024). PADRE tet^+^ CD4 T cells were positioned along either the trunk or one of the two possible branches of the trajectory (**Figure 2F; Supplementary** Figure 2B). Wishbone identified PSGL1^+^ T-bet^+^ SLAM^+^ Th1 cells as branch 1 and CXCR5^+^ PD-1^+^ BCL6^+^ TCF1^+^ FR4^+^ Tfh cells as branch 2 (**Figure 2C-F**). In contrast, the trunk of the Wishbone trajectory was composed of CD62L^hi^ CD44^low^ naive cells and CD62L^hi^ CD44^hi^ activated cells (**Figure 2B, F**). The CD44^hi^ CD62L^hi^ cells expressed high levels of TCF1 and SLAMF6 (**Figure 2C**), which are considered markers of stem-like CD8 T cells and stem-like Th1 precursors (Busselaar et al., 2020; Cardenas et al., 2024; Zou et al., 2024), thus suggesting the potential of CD44^hi^ CD62L^hi^ cells for further differentiation. In addition, CD44^hi^ CD62L^hi^ cells in the trunk of the trajectory had higher levels of CXCR5, BCL6 and PD-1 compared to naive cells, but still expressed PSGL1 (**Figure 2D, E**). Apart from being highly expressed by Tfh cells, PD-1 is also upregulated as a result of TCR triggering (Agata et al., 1996; Baumeister et al., 2016), and in line with this, all CD44^hi^ cells had higher expression of PD-1 compared to naïve, CD44^lo^ cells (**Figure 2E**). We also show that the pseudotime calculated by Wishbone matched real-time differentiation, as visualized by the decrease in frequency of CD62L^hi^ CD44^low^ and CD62L^hi^ CD44^hi^ cells over time (**Supplementary** Figure 2A**, C**). Altogether, these data indicate that, upon activation, antigen-specific CD4 T cells pass through a common CD44^hi^ CD62L^hi^ cell state before branching into Th1 or Tfh differentiation states.

### Th1 and Tfh cells develop from common precursor clones according to single-cell TCR sequencing

To characterize CD4 T-cell differentiation trajectories in an unbiased manner, we performed scRNAseq and paired TCRseq. For this purpose, mice received the MHC-II vaccine and CD3^+^ CD44^hi^ CD62L^low^ T cells were flow cytometrically purified from the dLN at days 5 or 10 after vaccination (**Supplementary** Figure 3A). After quality control filtering, selection of cells co-expressing TCRα and TCRβ sequences and exclusion of CD8 T cells and γδT cells (**Supplementary** Figure 3B), we obtained 4656 CD4 T cells for data analysis. Dimensionality reduction and clustering of cells based on their gene expression profiles identified 7 clusters that contained more than 20 cells (**Figure 3A, B; Supplementary** Figure 3C). Clusters 0 and 4 identified two Treg cell populations, characterized by expression of *Foxp3*, *Ctla4* and *Il2ra* (encoding CD25) (**Figure 3C**). Cluster 1 identified Tfh cells expressing *Tcf7* (encoding TCF1), *Cxcr5*, *Bcl6* and *Pdcd1* (encoding PD-1); cluster 2 identified Th1 cells expressing *Selplg* (encoding PSGL1), *Tbx21* (encoding T-bet), *Id2*, and *Cxcr3*; cluster 3 identified naive-like cells expressing *Ccr7*, *Lef1*, *Sell* (encoding CD62L), and *Tcf7*; whereas cluster 6 was dominated by an interferon (IFN)-stimulated gene (ISG) signature (**Figure 3C**). Interestingly, cluster 5 contained cells expressing both Th1-related genes (*Tbx21*, *Cxcr3*) and Tfh-related genes (*Tcf7*, *Cxcr5*, *Bcl6*, *Pdcd1*) (**Figure 3C**). The expression levels of these genes were intermediate between those found in Th1 cells (cl. 2) and Tfh cells (cl. 1). Cells of cluster 5 also expressed *Pdcd1* and *Cd69*, which are also markers of recent TCR triggering (**Figure 3C**) (Agata et al., 1996; Baumeister et al., 2016; Cibrián & Sánchez-Madrid, 2017; Simms & Ellis, 1996). Cluster 5 cells were higher in frequency at day 5 after vaccination than at day 10, supporting that they represent recently activated cells (**Figure 3B**). The frequency of Th1 cells (cl. 2) and Tfh cells (cl. 1) on the other hand, increased over time (**Figure 3B**), which aligns with ongoing effector differentiation of the activated CD4 T-cell pool.

**Figure 3:**
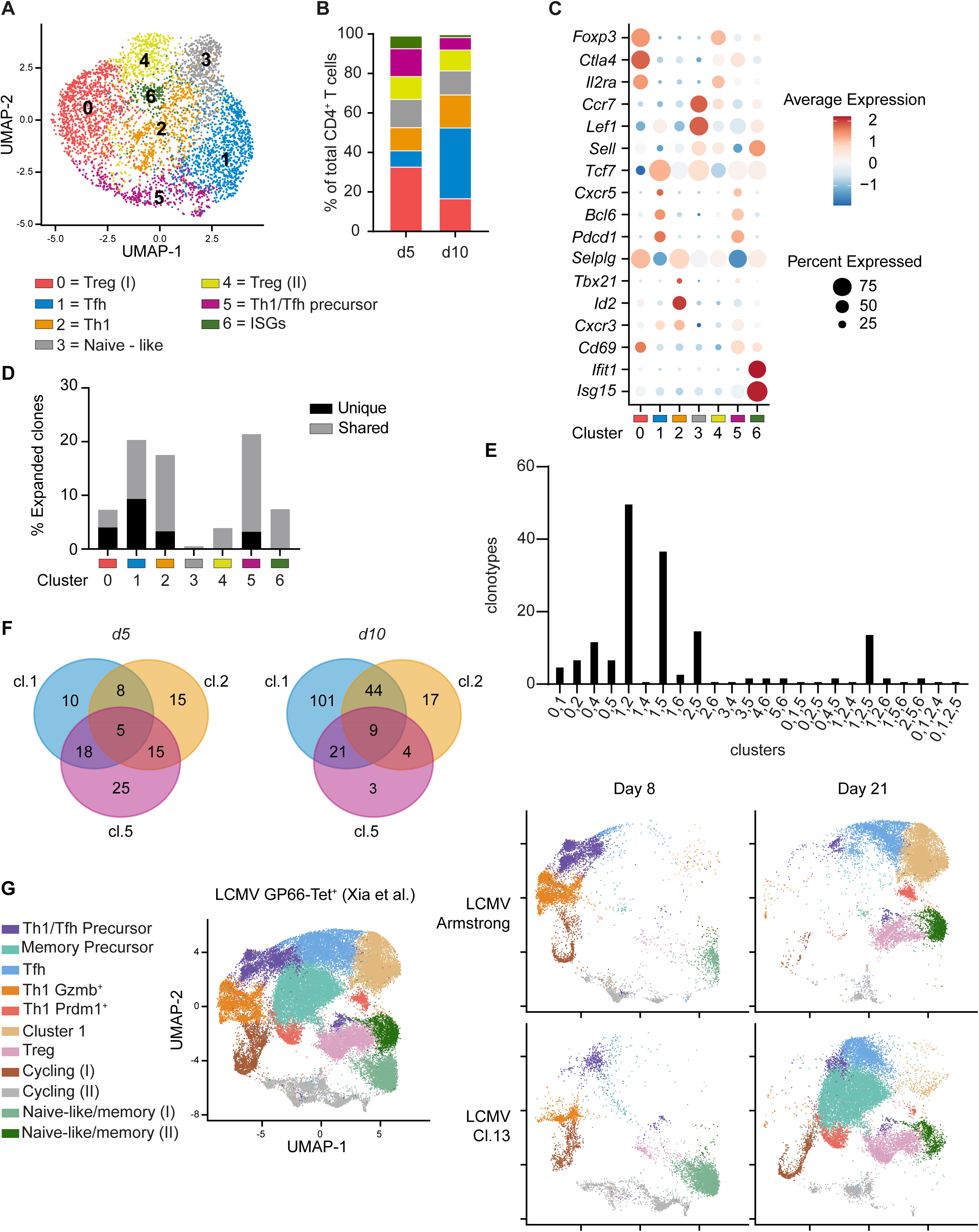
Identification of a common precursor of Th1 and Tfh cells. A) UMAP projection of CD4 T-cell clusters identified by scRNAseq of CD3^+^ CD44^+^ CD62L^-^ T cells isolated from the dLN of mice at days 5 and 10 after vaccination with the MHC-II plasmid. B) Bar plot depicting cluster distribution per time point. C) Average mRNA expression level of indicated differentiation-associated genes and percentage of positive cells per cluster. D) Frequency of expanded TCR clones (>1 cell per clone) per cluster. Black: TCR clones unique to one cluster; grey: TCR clones shared between two or more clusters. E) Quantification of clonotypes shared between two or more indicated clusters. F) Venn diagram depicting TCR sharing between indicated numbers of Tfh cells from cluster (cl.) 1, Th1 cells from cluster 2 and precursor cells from cluster 5 derived from dLN at day 5 (left) or day 10 (right) after vaccination. G) UMAP projection of LCMV-specific splenic CD4 T-cell clusters derived from LCMV-Armstrong and LCMV-clone 13 at day 8 and day 21 post infection from (Xia et al., 2022). Total cells (left). Deconvolution of UMAPs per infection type and day post infection is displayed on the right.

Next, to define clonal relationships between CD4 responder T cells in different states of effector differentiation, we paired the gene expression profiles per cell with TCRα and TCRβ sequence information. TCR clones were defined based on identical TCRα and TCRβ complementarity region (CDR)3 sequences, which identified a total number of 3549 clones within the CD4 T-cell compartment. We noted that Tfh cluster 1, Th1 cluster 2 and the intermediate phenotype cluster 5 had the highest frequency of expanded clones (>1 cell per clone) among all clusters (**Figure 3D)**. Additionally, the highest number of expanded clones was observed in Tfh cluster 1, followed by Th1 cluster 2, intermediate phenotype cluster 5 and Treg cluster 1 (**Supplementary** Figure 3D**, E**). Out of 352 expanded clones, 171 clones were shared between two or more clusters, while the rest was uniquely expanded within one cluster (**Supplementary** Figure 3D). Tregs of cluster 0 comprised more unique than shared clones and sharing occurred primarily with Tregs of cluster 4 (**Figure 3E, Supplementary** Figure 3D). The majority (72%) of expanded clones was shared between Tfh and Th1 cells (cl. 1 and 2), Tfh or Th1 cells and intermediate phenotype cells (cl. 5), or all three populations (**Figure 3E**). Accordingly, at days 5 and 10 after vaccination, TCR clonotype sharing between cluster 5 cells and Th1 cells, Tfh cells, or both was evident (**Figure 3F**). These data demonstrate the clonal relationship between cluster 5 cells and Th1 and Tfh cells, and confirm the indication of Wishbone analysis (**Figure 2F**) that antigen-activated CD4 T cells acquire an intermediate differentiation state expressing both Th1 and Tfh-related genes from which they can further develop into committed Th1 or Tfh cells. Cluster 5 cells thus represent the common precursors of Th1 and Tfh lineages. To determine whether Th1/Tfh precursor cells were formed not only after vaccination but also after infection, we used a published dataset of polyclonal GP66-tetramer^+^ antigen-specific CD4 T cells from mice infected with either acute LCMV-Armstrong or chronic LCMV-clone 13 (Xia et al., 2022) (**Figure 3G**). We aligned the transcriptional profile of Tfh (cluster 1), Th1 (cluster 2) and Th1/Tfh precursor cells (cluster 5) as defined in our vaccination setting with this dataset using module scoring. After LCMV infection, Tfh and two Th1 clusters were detected (**Supplementary** Figure 4A**, B**), with Th1 cells being prominent at day 8 in both infection models, and Tfh cells at day 21 (**Figure 3G**). The two Th1 clusters expressed classical Th1-associated genes including *Cxcr6, Tbx21, Id2,* and *Prf1,* but were distinguished by expression of either *Gzma* and *Gzmb* (Th1 Gzmb^+^ cluster) or *Prdm1, Eomes*, and *Gzmk* (Th1 Prdm1^+^ cluster) (**Supplementary** Figure 4E). The Tfh cluster expressed characteristic genes including *Cxcr5, Pdcd1, Il21 and Bcl6* (**Supplementary** Figure 4E). Importantly, at day 8 post infection with both acute and chronic LCMV, a population was enriched that matched the Th1/Tfh precursor cluster from our vaccination setting (**Figure 3G, Supplementary** Figure 4C). These cells expressed canonical Th1- and Tfh-associated genes (*Cxcr3*, *Cxcr5, Tbx21, Bcl6*) alongside key markers of the activated memory-precursor cell population described by (Xia et al., 2022) (*Tcf7, Slamf6, Batf, Zbtb32, Mif, Srm, Eif5a, Ncl, Hspd1*) (**Supplementary** Figure 4E). We also identified the previously described *Slamf6^+^ Tcf7^+^ Bcl6^lo^* memory-precursor cluster (Cardenas et al., 2024; Xia et al., 2022) to be specifically enriched at day 21 post-chronic LCMV infection. Despite the difference in their expression kinetics, the memory-precursor cluster shared features with the Th1/Tfh precursor cluster within the same dataset (**Figure 3G, Supplementary** Figure 4D**, 4E**). Altogether, these findings demonstrate that both after vaccination and virus infection, CD4 T cells transit through a common, uncommitted Th1/Tfh precursor state before diverging into Th1 and Tfh cells.

### Th1/Tfh precursors are proliferative and co-express Th1- and Tfh-related transcription factors and chemokine receptors

To gain deeper insights into the transcriptional programs underlying Th1 and Tfh cell differentiation, we performed differential gene expression analysis between Tfh (cl.1) and Th1 (cl.2) cells generated after vaccination. Gene signatures of the two T-helper subsets were characterized by lineage defining transcription factors, including *Tbx21*, *Id2*, *Bhlhe40* and *Prdm1* for Th1 cells, and *Bcl6*, *Tox* and *Tox2* for Tfh cells, which validated the robustness of our dataset (**Figure 4A**) (Nguyen et al., 2024; Oestreich et al., 2012; Shaw et al., 2016; Xu et al., 2019). Comparison of the transcriptional profile of the common Th1/Tfh precursor cells (cl. 5) to that of naive-like cells (cl. 3) revealed the characteristics of this population (**Figure 4B**). Cluster 5 cells expressed many genes associated with either Th1 or Tfh fate, including transcription factors *Tbx21*, *Bhlhe40*, *Prdm1* and *Bcl6*, as well as chemokine receptors *Cxcr3* and *Cxcr5*. These transcripts were co-expressed within the same cells and enriched in cluster 5 at day 5 post-vaccination, as shown by the positive correlation in expression of e.g. *Tbx21* and *Cxcr5*, or *Tbx21* and *Bcl6* (**Supplementary** Figure 5A**, B**). In addition, cells of cluster 5 displayed upregulation of *Mki67*, indicating active cell division, and multiple costimulatory receptors including *Icos*, *Tnfrsf8* (encoding CD30)*, Tnfrsf9* (encoding CD137, alias 4-1BB), *Tnfrsf18* (encoding CD357, alias GITR), and *Tnfrsf1b* (encoding CD120b, alias TNFR2) (**Figure 4B)**. They also gained expression of the metabolic enzyme lactate dehydrogenase (*Ldha*) (**Figure 4B**), which has been associated with the formation of stem-like Th1 precursors (Zou et al., 2024).

**Figure 4:**
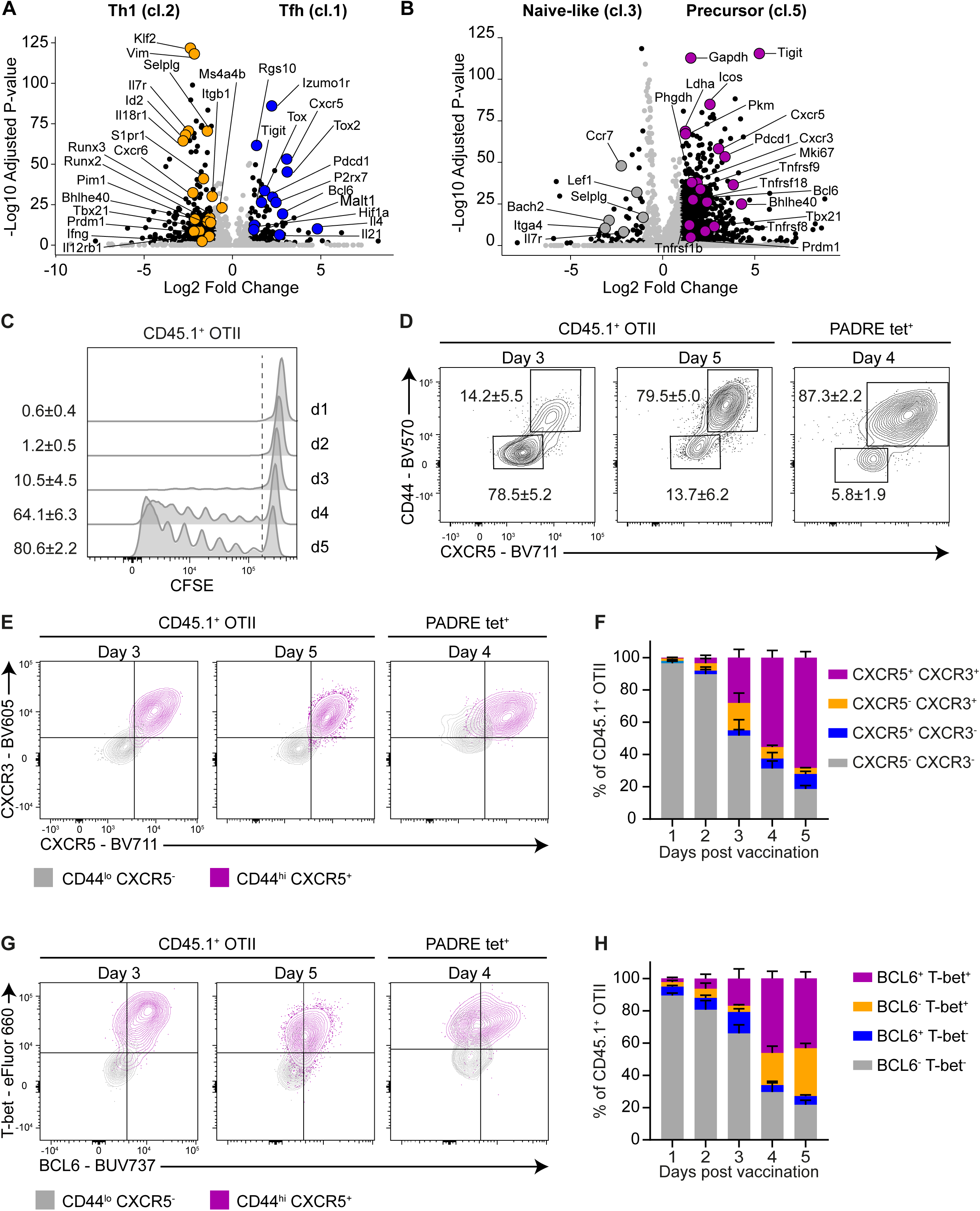
Common Th1/Tfh precursors co-express Th1- and Tfh markers at protein level. A, B) Volcano plots depicting differential gene expression derived from scRNAseq data, comparing Tfh cells (cl. 1) versus Th1 cells (cl. 2) (A), or naive-like CD4 T cells (cl. 3) versus the common Th1/Tfh precursor cells (cl. 5) (B). Black dots depict genes with fold change < -1 or > 1 and adjusted P<0.05. C) Mice received OTII cells and MHC-II:OVA vaccine and dLNs were harvested and cells analyzed by flow cytometry. Histograms depicting divisions of vaccine-activated, adoptively transferred OTII cells, as measured by CFSE dilution. Numbers indicate frequency of divided OTII cells (mean ± SD). D) Representative CD44 and CXCR5 protein expression on CD45.1^+^ OTII or PADRE tet^+^ CD4 T cells, measured in dLNs at indicated days post-vaccination. E, G) Representative overlays of CXCR5 and CXCR3 (E) or T-bet and BCL6 (G) protein expression in CD44^lo^ CXCR5^-^ (gray) and CD44^+^ CXCR5^+^ (purple) CD45.1^+^ OTII or PADRE tet^+^ CD4 T cells. F, H) Proportion of CD45.1^+^ OTII responder cells expressing CXCR5 and CXCR3 (F) or T-bet and BCL6 (H) in the dLNs at the indicated days post-vaccination (N=3-5 per time point; representative of 3 independent experiments). Graphs depict mean ± SD.

To map the formation and function of common Th1/Tfh precursor cells by flow cytometry, we followed the response of TCR transgenic OTII cells that recognize OVA_323-339_ peptide in the context of MHC-II. CD45.1^+^ OTII cells were labeled with carboxyfluorescein succinimidyl ester (CFSE) to track proliferation and adoptively transferred into CD45.2^+^ recipient mice. One day later, recipient mice received a modified MHC-II DNA vaccine encoding both PADRE and OVA_323-339_ epitopes (MHC-II:OVA; **Supplementary** Figure 5C). This vaccine simultaneously induced an OTII T-cell response to the OVA epitope and an endogenous CD4 T-cell response to the PADRE epitope (**Supplementary** Figure 5D**, E**), enabling us to compare the differentiation trajectories of these two responder cell populations. Because the transferred naïve OTII cells have a relatively high frequency, this system allowed us to analyze their early activation state at day 3 post-vaccination. Endogenous PADRE tet^+^ CD4 T cells became detectable by tetramer staining at day 4 post-vaccination (**Supplementary** Figure 5F**, G**) and we therefore monitored the CD4 T-cell response in the dLNs from day 1 to 5 after vaccination. At day 3, OTII cell proliferation was initiated and at day 4, the majority of OTII cells were actively dividing, with a proportion having undergone at least seven divisions already (**Figure 4C, Supplementary** Figure 5H). Accordingly, the frequencies of both OTII cells and PADRE tet^+^ CD4 T cells were significantly increased by day 4 post-vaccination (**Supplementary** Figure 5F**, G**). At day 3, concurrent with the start of cell division, OTII cells upregulated the activation marker CD44, as well as the chemokine receptor CXCR5 (**Figure 4D, Supplementary** Fig 5I) that is commonly used as a marker of Tfh precursors (Baumjohann et al., 2011; Y. S. Choi et al., 2011; Ma et al., 2012). Interestingly, CD44^hi^ CXCR5^+^ OTII cells also expressed high levels of CXCR3 (**Figure 4E, F**), a chemokine receptor that among CD4 T cells is thought to be specifically upregulated on Th1 precursors (Ditoro et al., 2018). Consistent with our scRNAseq data, CD44^hi^ CXCR5^+^ CXCR3^+^ OTII cells co-expressed T-bet and BCL6 proteins (**Figure 4G, H**). Within the endogenous CD4 T-cell pool, PADRE tet^+^ CD44^hi^

CD4 T cells also co-expressed CXCR5, CXCR3, T-bet and BCL6 at day 4 post-vaccination (**Figure 4E, G; Supplementary** Figure 5J-L). Furthermore, CD44^hi^ OTII cells expressed significantly higher levels of PD-1 compared to CD44^lo^ cells (**Supplementary figure 5M**), consistent with our results for endogenous cells (**Figure 2E**; **Figure 3C**). TCR transgenic T cells all have the same affinity and may therefore have an intrinsic predisposition to a specific differentiation trajectory (Künzli & Masopust, 2023). Next to low affinity OTII cells, we therefore also used SMARTA T cells that have a high affinity TCR recognizing the GP_61-80_ epitope of LCMV in the context of MHC-II (Ditoro et al., 2018) in conjunction with a modified MHC-II vaccine that encodes both the PADRE and GP_61-80_ epitopes (**Supplemental Figure 5N and O**). SMARTA CD4 T cells also differentiated into CD44^+^ precursor cells co-expressing CXCR3, CXCR5, T-bet and BCL6, as observed at day 4 post-vaccination (**Supplemental Figure 5O**). Obtaining the same results with low and high affinity TCR transgenic CD4 T cells, as well as PADRE-specific endogenous, polyclonal CD4 T cells indicates that the formation of the Th1/Tfh precursor cell pool of this specific phenotype is independent of TCR affinity. The collective data establish that activated CD4 T cells initially form a proliferative pool of bipotent Th1/Tfh precursors with a distinct marker profile that can differentiate into either Th1 or Tfh cells, independent of TCR affinity.

### Specific costimulatory signals driving formation of the common Th1/Tfh precursor pool and subsequent Th1- or Tfh differentiation

Having identified the common Th1/Tfh precursor, we were interested in the costimulatory signals that drive its formation and further differentiation. It is known that Th1 and Tfh differentiation rely on specific costimulatory signals (Schorer et al., 2019), but the impact of these has not previously been mapped in the differentiation trajectory that we have established here. We first examined the impact of costimulatory receptor CD28 that synergizes with the TCR/CD3 complex in initiating T-cell activation, but reportedly is also required to maintain both Th1 and Tfh cells (Linterman et al., 2014). Mice received OTII cells and MHC-II:OVA-based vaccination and CD28 costimulation was inhibited by infusion of blocking monoclonal antibodies (mAbs) to both CD80 and CD86. This intervention did not significantly affect the overall frequency of OTII cells in dLN at day 5 after vaccination (**Supplementary** Figure 6A), but did reduce the frequency of CD44^+^ CXCR3^+^ CXCR5^+^ OTII Th1/Tfh precursor cells (**Figure 5A**). CD80 and CD86 blockade also reduced the frequency of total PADRE tet^+^ CD4 T cells (**Figure 5B**), thus showing that CD28 costimulation promotes formation of the Th1/Tfh precursor pool by supporting clonal expansion. Because CD80 and CD86 have different affinities for CD28 (Collins et al., 2002) and distinct roles in the maturation of Tfh cells in GCs (Wang et al., 2015), we also investigated whether they had independent impact on the CD4 T-cell differentiation trajectory. Single blockade of either CD80 or CD86 did not affect the generation of CXCR3^+^ CXCR5^+^ OTII Th1/Tfh precursor cells (**Figure 5C**), nor the expansion and initial Th1 or Tfh bifurcation of PADRE^+^ CD4 T cells (**Figure 5D; Supplementary** Figure 6B-D), indicating that CD80 and CD86 are redundant in promoting formation of the Th1/Tfh precursor pool.

**Figure 5:**
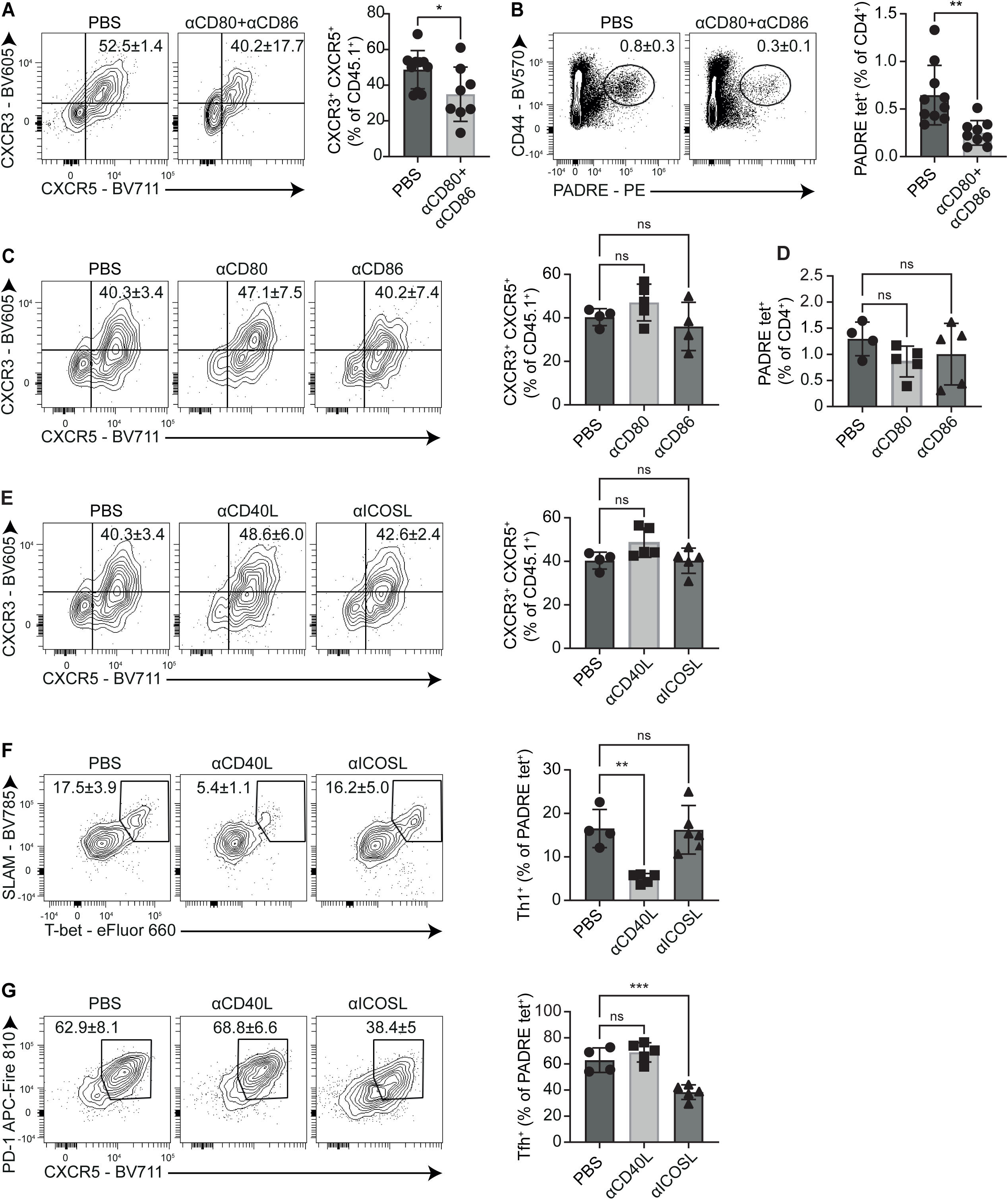
CD28 costimulation promotes formation of the Th1/Tfh precursor pool, from which CD40 or ICOS drive Th1 or Tfh differentiation, respectively. Mice received OTII cells and MHC-II:OVA vaccine and were treated with blocking mAbs to indicated costimulatory ligands or PBS (control). At day 5 after vaccination, cells from dLNs were analyzed by flow cytometry. Representative plots and quantification of frequencies of cells with indicated markers are shown. (A, B) Frequencies of CXCR3^+^ CXCR5^+^ CD45.1^+^ OTII cells (A) and PADRE tet^+^ CD4 T cells in mice treated with a combination of blocking mAbs to CD80 and CD86 (B) (N=9-10 per group; data pooled from 2 independent experiments). Unpaired student’s-t test. C, D) Frequencies of CXCR3^+^ CXCR5^+^ CD45.1^+^ OTII cells (C) and PADRE tet^+^ CD4 T cells (D) in mice treated with blocking mAbs to either CD80 or CD86. E) Frequencies of CXCR3^+^ CXCR5^+^ precursor CD45.1^+^ OTII T cells in mice treated with blocking mAb to either CD40L or ICOSL F, G) Frequencies of SLAM^+^ T-bet^+^ Th1 (F) and CXCR5^+^ PD-1^+^ Tfh (G) PADRE tet^+^ CD4 T cells in mice treated with blocking mAb to either CD40L or ICOSL. C-G) N=4-5 per group; data are representative at least two independent experiments. One-Way ANOVA with Dunnett post hoc test was used to correct for multiple testing when comparing inhibitory conditions to control. All graphs depict mean ± SD.

We next evaluated the impact of CD40 and ICOS costimulation on the CD4 T-cell differentiation trajectory. CD40:CD40L interaction has been reported to promote both Th1 and Tfh function (Baumjohann et al., 2013; Grewal et al., 1995; Liu et al., 2014) and ICOS:ICOSL interaction is important for Tfh differentiation (Baumjohann et al., 2013; Y. S. Choi et al., 2011; Liu et al., 2014; Shin et al., 2015). Blockade of CD40L or ICOSL did not affect the frequency of CD44^+^ CXCR5^+^ CXCR3^+^ OTII precursor cells, total PADRE tet^+^ CD4 T cells, or total OTII cells (**Figure 5E; Supplementary** Figure 6E-F), indicating that CD40- and ICOS costimulation were not required for formation of the Th1/Tfh precursor pool. However, among PADRE tet^+^ cells, CD40L blockade significantly reduced the frequency of Th1 cells, while not affecting the frequency of Tfh cells (**Figure 5F-G**). Conversely, blockade of ICOSL lowered the frequency of Tfh cells, while not affecting the frequency of Th1 cells (**Figure 5F, G**). Altogether, these data provide a scenario wherein CD28 costimulation driven by either CD80 or CD86 promotes formation of the Th1/Tfh precursor cell pool, whereafter the precursors are driven into Th1 cells with the aid of CD40-CD40L interactions and into Tfh cells with the aid of ICOS-ICOSL interactions.

### Th1/Tfh precursors differentiate into either Th1 or Tfh cells in a second priming step on cDC1s or B cells, respectively

We next examined the role of cDC1s and B cells in guiding the differentiation of Th1 and Tfh cells from the Th1/Tfh precursor. The role of cDC1s was determined by using *Batf3*^-/-^ mice on a C57BL6/J background (Edelson et al., 2010, 2011; Hildner et al., 2008), which display complete loss of migratory cDC1s (MHCII^hi^ CD11c^int^ CD103^+^ XCR1^+^) and partial loss of resident cDC1s (MHCII^int^ CD11c^hi^ XCR1^+^) (**Supplementary** Figure 7A-F). The role of B cells was determined by effective B-cell depletion using a mAb to CD20 prior to vaccination (**Supplementary** Figure 7G). In both experimental set-ups, OTII cells were adoptively transferred, the MHC-II:OVA vaccine was administered, and responses were read out in the dLNs by flow cytometry at day 5 after vaccination. Neither *Batf3*-deficiency (**Supplementary** Figure 7H**, I**), nor B cell depletion (**Supplementary** Figure 7L**, M**) affected the clonal expansion of endogenous PADRE tet^+^ CD4 T cells or OTII cells. Formation of CD44^+^ CXCR3^+^ CXCR5^+^ Th1/Tfh precursor cells was also unaffected in mice lacking these APCs (**Figure 6A, B**). However, *Batf3*^-/-^ mice showed a decrease in the generation of Th1, but not Tfh cells, while B-cell depleted mice showed a decrease in the generation of Tfh, but not Th1 cells. This was the case for both endogenous PADRE tet^+^ CD4 T cells (**Figure 6C-F**) and for exogenous OTII cells (**Supplementary** Figure 7J-K**, N-O**). Altogether, these data suggest that CD28-dependent formation of the Th1/Tfh precursor pool is instructed by APCs other than cDC1s and B cells, most likely cDC2s. Thereafter, in a second step of priming, the precursor cells are instructed for Th1 differentiation on cDC1 as driven by CD40 costimulation, or for Tfh differentiation on B cells, as driven by ICOS costimulation (**Figure 6G**).

**Figure 6:**
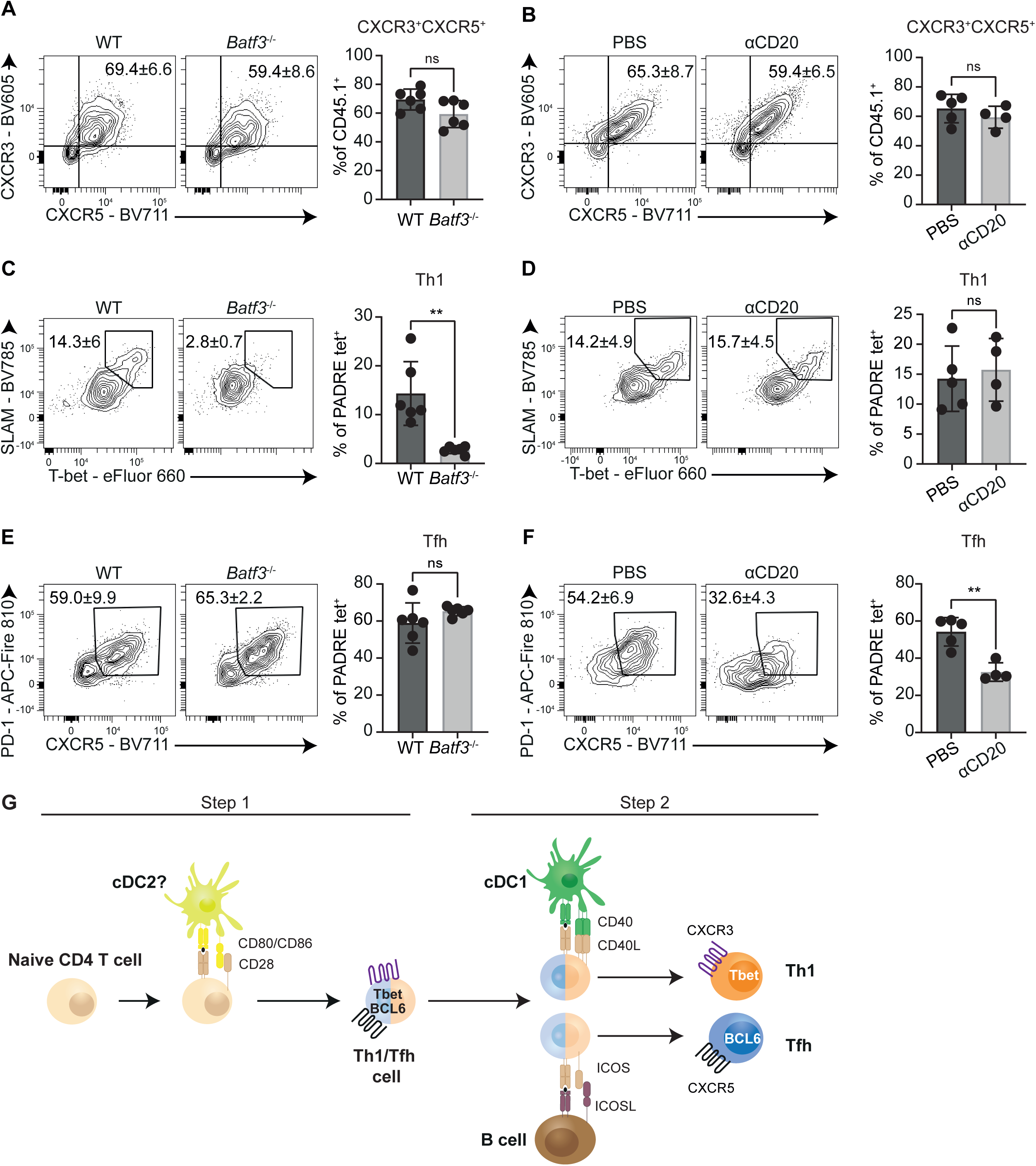
Th1 or Tfh differentiation from the common Th1/Tfh precursor is driven by interactions with cDC1s or B cells, respectively. Wild-type (WT) or Batf3-/- mice received OTII cells and MHC-II:OVA vaccine and were treated with depleting mAb to CD20 or control (PBS) as indicated. At day 5 after vaccination, cells from dLN were analyzed by flow cytometry. Representative plots and quantification of frequencies of cells with indicated markers are shown. (A, B) Flow cytometry plots and quantification of CXCR3^+^ CXCR5^+^ CD45.1^+^ OTII cells in the dLN of WT versus *Batf3^-/^*^-^ mice (A), or WT mice that were treated with B-cell depleting anti-CD20 mAb or PBS (control), two days before vaccination (B). C-F) Flow cytometry plots and quantification of SLAM^+^T-bet^+^ Th1 (C, D) or CXCR5^+^ PD-1^+^ Tfh (E, F) PADRE tet^+^ CD4 T cells in WT versus *Batf3^-/-^* mice (C, E), or WT mice treated with anti-CD20 mAb or PBS (D, F). All graphs depict mean ± SD (N=6 per group in A, C, E; N=4-5 per group in B, D, F). A-F, Unpaired student’s t-test. G) Proposed model of cellular and molecular interactions that, next to TCR/CD3 signals, drive CD4 T-cell differentiation towards Th1 or Tfh cells.

## Discussion

In this study, we show that upon *in vivo* activation, individual clones within the endogenous CD4 T-cell repertoire transit through a bipotent Th1/Tfh precursor state before committing to Th1 or Tfh cell differentiation. These precursor cells are in an early stem-like differentiation state characterized by TCF1 and SLAMF6, and they co-express Th1- and Tfh-related markers at mRNA and protein level. Th1/Tfh precursor cells retain the capacity to differentiate into both Th1 and Tfh cells, with TCR sharing underscoring their common origin. They arise following vaccination as well as acute or chronic LCMV infection. Th1/Tfh precursor cells resemble, but are distinct from, previously described memory-precursor populations, which are specifically enriched upon chronic LCMV infection (Xia et al., 2022). Instead, they align with a previous study identifying a common Th1/Tfh precursor among *Plasmodium chabaudi*-specific TCR transgenic CD4 T cells (Lönnberg et al., 2017). Together, these findings establish the Th1/Tfh precursor state as a common feature in the CD4 T-cell differentiation trajectory in type 1 immunity.

During priming in secondary lymphoid organs, T cells have multiple contacts with different APCs. This is most explicit for Tfh formation in which both cDCs and B cells play a role. CD4 T cells are generally primed by cDC2s in the T-cell zone (Gerner et al., 2017; Krishnaswamy et al., 2017; Li et al., 2016), giving rise to CXCR5-expressing cells that are generally referred to as pre-Tfh cells. These cells migrate along a CXCL13 gradient from the T-cell zone towards the border of the B-cell follicle or the interfollicular region (IFR) (Crotty, 2014; Kerfoot et al., 2011). At that location, they receive further stimulation to enter the B-cell follicle and complete Tfh differentiation (Kerfoot et al., 2011; Kim et al., 2024; Song & Craft, 2024). Although expression of CXCR5 and PD-1 on activated CD4 T cells early after vaccination is generally considered indicative of Tfh commitment, not all CXCR5^+^ PD-1^+^ CD4 T cells become Tfh cells (Kerfoot et al., 2011; Song & Craft, 2024). This can now be explained by our definition of the CXCR3^+^ CXCR5^+^ Th1/Tfh precursor state. Thus far, CXCR3 that binds IFN-responsive chemokines CXCL9 and CXCL10, is generally associated with Th1 differentiation (Groom et al., 2012). This chemokine receptor promotes migration of activated CD4 T cells from the T-cell zone towards the IFR of the LN (Groom et al., 2012). Here, CD4 T-cell help for CD8 effector and memory differentiation is likely delivered upon cognate contact of both CD4 and CD8 T cells with the same cDC1 (Duckworth et al., 2021; Eickhoff et al., 2015; Hor et al., 2015). However, activated CD4 T cells can also interact with B cells in the IFR (Kerfoot et al., 2011). In line with the existence of the Th1/Tfh precursor, CXCR3 has previously been found on cells with a Tfh phenotype (Künzli & Masopust, 2023; Weinstein et al., 2018). We show that CXCR3^+^ CXCR5^+^ cells are bipotent Th1/Tfh precursors and propose that they correspond to previously defined pre-Tfh cells. We postulate that the differentiation fate of these cells depends on the dominance of either of the chemokine cues, allowing CXCR5-mediated guidance to a B cell for Tfh differentiation or CXCR3-mediated guidance to a cDC1 for Th1 differentiation.

Accordingly, we found that cDC1 and CD40:CD40L interaction are required for Th1 differentiation from the common precursor and B cells and ICOS costimulation are required for Tfh differentiation. CD40 signaling in cDC1 is associated with upregulation of costimulatory molecules such as CD70, expression of IL-12 and apoptosis resistance (Lei et al., 2024; Schulz et al., 2000; Wu et al., 2022). CD40 is also critical in the maintenance of Tfh and GC responses (Baumjohann et al., 2013), and it was somewhat surprising that blockade did not affect Tfh cell differentiation in our model. Similarly, CD86 is known to promote Tfh differentiation (Wang et al., 2015), but we did not find any effects of single CD86 blockade. This may be explained by the fact that we focused on early differentiation stages, prior to the establishment of germinal centers, when CD40 and CD86 signals may be trivial or compensated for. In addition to cDC1, CD40 signals and CXCR3-attracting chemokines can also be delivered by monocyte-derived (Mo)DCs present within the IFR (Duckworth et al., 2021). This aligns with our recent data showing that CD4 T cells and MoDCs communicate in a CD40-dependent feed-forward loop to enforce Th1 effector differentiation (Bosma et al., 2025), which can be mediated by IL-12 (De Koker et al., 2017; Leal et al., 2021).

The formation of the bipotent Th1/Tfh state allows immune system flexibility in tailoring host protection to ongoing infections. Different viral infections can result in different Th1/Tfh ratios, which are reinforced through regulatory feedback loops that suppress the expression of T-bet or BCL6 (J. Choi et al., 2020; De Giovanni et al., 2020; Duckworth & Groom, 2021; Sheikh & Groom, 2021). Indeed, LCMV-Armstrong and Influenza infection differentially depend on T-bet expression (Sheikh & Groom, 2021). Similarly, vesicular stomatitis virus (VSV) induces a higher Tfh/Th1 ratio than LCMV by inducing an early wave of IFN-I and IL-6 (De Giovanni et al., 2020). Thus, the formation of Th1/Tfh precursors and their subsequent differentiation will be context dependent. Overcoming a certain threshold of T-bet versus BCL6 expression is likely essential to consolidate Th1 or Tfh differentiation (Oestreich et al., 2012).

In human peripheral blood, CXCR3^+^ CXCR5^+^ PD-1^lo^ cells have been found, as well as Tfh-like cells that share chemokine receptor and transcription factor expression with Th2 or Th17 cells (Morita et al., 2011). CXCR3^+^ Tfh-like cells have been associated with improved responses to influenza vaccination and correlate with neutralizing antibodies in HIV controllers (Bentebibel et al., 2013, 2016; Martin-Gayo et al., 2017). These circulating Tfh-like cells express CD62L and CCR7, suggesting that they are in a less differentiated or memory-like state (Morita et al., 2011) and remain responsive to antigenic stimulation. Possibly, these cells represent the Th1/Tfh precursors we have identified here.

The stem-like CD4 T cells that have been identified in transplantation rejection and cancer, and precede Th1 differentiation (Cardenas et al., 2024; Zou et al., 2024), may also represent Th1/Tfh precursors. In transplantation models, these stem-like CD4 T cells relied on metabolic rewiring, including *Ldha* expression (Zou et al., 2024), which was also upregulated in Th1/Tfh precursors following vaccination. In cancer models, they differentiated into Th1 cells upon Treg depletion (Cardenas et al., 2024). These stem-like CD4 T cells have primarily been characterized using markers that are established for CD8 T cells. Many of these markers, including TCF1, PD-1 and SLAMF6, are also expressed by the common Th1/Tfh precursor, supporting the general concept of a stem-like, highly proliferative and uncommitted cell state. A recent pan-cancer analysis across 21 tumor types further supports this concept, identifying a tumor-enriched CD4 T-cell population predicted to be tumor-specific and characterized by both Th1- and Tfh-associated gene expression (Zheng et al., 2021). Together, these findings underscore the need to generally acknowledge the Th1/Tfh precursor cell state and to fully define the developmental cues for either Th1 or Tfh differentiation from this state.

## Materials and Methods

### Mice

Mice were housed in individually ventilated cages (IVC) under specific pathogen free (SPF) conditions and with *ad libitum* access to food and water. Ambient temperature was ∼19-21°C and humidity 30-70%. Lighting was provided on a 12 h light/dark cycle. Eight-week-old male and female C57BL/6JRj mice were purchased from Janvier laboratories (Le Genest Saint Isle, France). OTII*CD45.1 ((TcraTcrb)425Cbn/CrlLumc) on a C57BL/6JRj background were bred in house. Batf3^-/-^ (B6.129S(C)-Batf3tm1Kmm/J) on a C57BL/6J background and wild type controls were obtained from The Jackson Laboratory. All mouse experiments were performed in accordance with institutional and national guidelines and were approved by the Animal Welfare Body at Leiden University Medical Center.

### DNA tattoo vaccination

The cDNAs encoding the MHC-II, MHC-II:OVA and control sequences used for DNA vaccination were expressed in a pVAX1 plasmid vector. MHC-II and control vaccines have previously been described (Ahrends et al., 2016; Oosterhuis et al., 2012). MHC-II:OVA and MHC-II-GP(61-80) vaccines were generated by excising the cassette encoding the MHC-II epitopes (p30, PADRE, HIV Nef_56-68_) from the MHC-II vaccine cDNA with HindIII and XhoI restriction enzymes, and replacing it with a gBlock containing the MHC-II cassette linked to nucleotide sequences encoding OVA_323-339_ (ISQAVHAAHAEINEAGR) or LCMV-Armstrong derived GP_61-80_ (GLKGPDIYKGVYQFKSVEFD) and a GPGPGPG linker on both ends as for the original MHC-II vaccine.

For intraepidermal DNA tattoo vaccination (Ahrends et al., 2016; Babała et al., 2018), mice were anesthetized using isoflurane (induction phase: 4% isoflurane, airflow 0.8 l/min; maintenance phase: 2% isoflurane, airflow 0.4 l/min) and the right hind leg of mice was depilated using Veet containing limonene (Reckitt Benckiser). Subsequently, 15 ml of a 2 mg/ml plasmid DNA in H_2_O was tattooed into the skin with a Permanent Make Up tattoo machine (Cheyenne, MT Derm GmbH) using a sterile disposable needle with a needle depth of 1 mm. Tattooing was performed for 45 sec, oscillating at a frequency of 100 Hertz. Mice were vaccinated at days 0, 3, and 6 unless otherwise indicated, and alternating between the internal or external upper part of the right hind leg.

### Adoptive T-cell transfer

For adoptive transfer, spleens were isolated from 8- to 12-week-old, sex matched OTII*CD45.1 mice after euthanasia. Spleens were mechanically disrupted and passed through a 70 mM Falcon strainer (Corning). Subsequently, single cell suspensions were incubated in red blood cell (RBC) lysis buffer (Santa Cruz) for 5 min at room temperature. CD4 T cells were isolated by negative selection using a CD4 T-cell isolation kit (Miltenyi) according to the manufacturer’s protocol. If indicated, purified OTII*CD45.1 CD4 T cells were labelled with 2.5 mM CFSE (Invitrogen) for 10 min at 37°C. Free CFSE was then quenched by adding at least five times excess of Iscove’s modified Dulbecco’s medium (IMDM) with L-glutamine (Capricorn) and 10% fetal bovine serum (FBS), 100 units/ml penicillin and 100 mg/ml streptomycin (Gibco), and resting cells for 5 min at room temperature. Cell purity (>90%) and CFSE-labelling were checked for each experiment by flow cytometry using the following monoclonal antibodies (mAbs) against mouse molecules: 1:100 CD45.1 BUV395 (clone A20, BD Biosciences), 1:200 CD4 BUV805 (clone GK1.5, BD Biosciences), 1:200 TCRva2 BV421 (clone B20.1),TCRvb5.1/5.2 (clone MR9-4), 1:100 CD44 BV570 (clone IM7), and 1:100 CD62L PE-Fire 810 (clone W18021D; all BioLegend). C57BL/6JRj mice received 0.1 x 10^6^ or 0.5 x 10^6^ or 1 x 10^6^ OTII*CD45.1 cells in 100 µl Hank’s Balanced Salt Solution (HBSS) depending on the day of read out, or 5 x 10^5^ purified SMARTA*GFP (B6.Cg-Ptprca Pepcb Tg(TcrLCMV)1Aox/PpmJx) cells in 100 µl HBSS, kindly provided by Dr. K. Jobin and Prof. W. Kastenmüller (University of Würzburg, Germany), by retro-orbital i.v. injection.

### Antibody and reagent treatment of mice

All antibodies or reagents were injected intraperitoneally (i.p.). Mice received 250 mg depleting anti-CD20 mAb (MB20-11, BioXcell) at day -2. Mice with incomplete B-cell depletion were excluded from analysis. For blocking costimulatory ligands, mice received injections on days 0 and 3 post vaccination with 150 mg anti-CD40L mAb (MR-1, BioXcell), 200 mg anti-CD80 mAb (1G10, BioXcell), 200 mg anti-CD86 mAb (GL-1, BioXcell) or 200 mg anti-ICOSL mAb (HK5.3). FTY720 (Fingolimod, Cayman Chemicals) was dissolved in 96% ethanol at 20 mg/ml and prior to injection diluted in 0.9% NaCl. Mice received 25 mg FTY720 in 200 ml 0.9% NaCl at days 0, 3 and 6.

### Cell isolation

On the day of readout, mice were euthanized and organs collected. Blood was collected by tail vein puncture or cardiac puncture. Erythrocytes were lysed by incubation in RBC lysis buffer (Santa Cruz) for 5 min at room temperature. Inguinal dLNs of the vaccination site were punched repeatedly with a 25G needle and incubated for 30 min at 37°C while shaking in 100 mg/ml Liberase TL (Roche) prepared in 500 ml DMEM. Enzymatic digestion was quenched by adding FBS-supplemented IMDM, and single cell suspensions prepared by mechanical disruption of dLNs through 70 mm Falcon strainers (Corning).

### Flow cytometry

For cell surface staining, mAbs were diluted in FACS buffer (PBS + 2% FBS). Single cell suspensions were first incubated for 1 h at room temperature with 1:200 PE-conjugated PADRE tetramer (AKFVAAWTLKAA in I-A(b), NIH tetramer facility), and then for 30 min on ice with the following mAbs: 1:200 CD4 BUV805 (clone GK1.5), 1:200 PSGL1 BUV661 (clone 2PH1), 1:200 CD8a BV750 (clone 53-6.7), 1:00 CD8a V500 (clone 53-6.7), 1:400 CD95 BV605 (clone Jo2), 1:100 CD45.1 BUV395 (clone A20, all BD Biosciences), 1:200 CD138 (clone 281-2), 1:400 GL7 FITC (clone GL-7), 1:600 IgD PerCP-Cy5.5 (clone 11-26c2a), 1:100 CD44 BV570 (clone IM7), 1:100 CD44 BV785 (clone IM7), 1:100 CD62L PE-Fire 810 (clone W18021D), 1:100 PD-1 APC-Fire 810 (29F.1A12), 1:100 SLAM BV785 (clone TC15-12F12.2), 1:100 SLAMF6 Pacific Blue (clone 330-AJ), 1:100 CD3 PE-Fire 700 (17A2), 1:50 CXCR5 BV711 (clone L138D7), 1:50 CXCR3 BV605 (clone CXCR3-173, all BioLegend), 1:200 TACI APC (clone ebio8F10-3, eBioscience), 1:800 CD19 BUV661 (clone ebio1D3, Invitrogen) and APC-conjugated E7 tetramer (E7(49-57) RAHYNIVTF in H-2D^b^, produced in house). Dead cells were excluded using 1:500 Zombie UV fixable viability dye (BioLegend) or 1:1000 LIVE/DEAD NIR fixable viability dye (Invitrogen).

For intracellular staining of T cell-related markers, cells were fixed with the Foxp3/Transcription factor staining kit (eBioscience). For intracellular staining of B cell-related markers, cells were fixed with BD Cytofix/Cytoperm (BD Biosciences). In both cases, fixation was performed for 30 min on ice. After fixation, cells were either permeabilized and intracellularly stained directly or washed with FACS buffer and frozen gradually at -80°C in 10% DMSO and 90% FBS. Upon thawing, cells were washed with FACS buffer and re-fixed for 5 min at room temperature prior to permeabilization and intracellular antibody staining. The following mAbs were used: 1:50 BCL6 BUV737 (clone K112-91), 1:50 GATA3 BUV395 (clone L50-823), 1:100 RORγt BV480 (clone Q31-378, all BD Biosciences), 1:100 TCF1 Alexa fluor 488 (clone C63D9, Cell Signalling Technology), 1:200 T-bet eFluor 660 (clone ebio4B10, eBioscience) and 1:100 FOXP3 PE-Cy5 (clone FJK16s, Invitrogen). Data were acquired using a Cytek Aurora Spectral Flow Cytometer equipped with 355 nm, 405 nm, 488 nm, 561 nm and 640 nm lasers (Cytek Biosciences), and analysed with OMIQ (Dotmatics version 2024.07) or FlowJo software (TreeStar, version 10.8.1).

UMAP analysis and Wishbone trajectory analysis were performed using OMIQ on a maximum of 1000 PADRE^+^ CD4^+^ T cells per mouse (N=5/time point, total 27867 cells). For dimension reduction and Wishbone analysis, the following markers were included: PSGL1, FR4, BCL6, SLAMF6, CD44, CXCR5, SLAM, TCF1, CD62L, T-bet and PD-1. For Wishbone analysis, diffusion maps were generated first and then Wishbone was performed from diffusion components, with naïve (CD62L^+^ CD44^-^ PD-1^-^ SLAMF6^-^) cells selected as starting point.

### Preparations for transcriptome analysis

Mice received MHC-II vaccine and cells were harvested at day 5 or 10 after the first vaccination. Single cell suspensions of dLNs from 3 mice per time point were pooled to reduce biological variability. Samples were incubated with 1:50 anti-mouse CD16/32 mAb (Fc Block^TM^, clone 2.4G2, BD Biosciences) in BD Stain buffer (with FBS, BD Biosciences) for 5 min on ice and subsequently stained with the following mAbs in BD stain buffer: 1:100 CD3 BV421 (clone 17A2, Biolegend), 1:100 CD19 PerCP-Vio700 (clone REA749, Miltenyi Biotech), 1:100 CD44 PE-Cy7 (clone IM7, eBioscience), 1:100 CD62L FITC (clone Mel-14, eBioscience) and 1:1000 LIVE/DEAD Fixable Near-IR dye (Invitrogen). CD3^+^ CD44^hi^ CD62L^lo^ cells were FACS-sorted on a FACSAria II (BD Biosciences). Subsequently, samples were stained with unique hashtag oligonucleotide (HTO)-conjugated antibodies to CD45/MHC-I (TotalSeq C hashtags, C0301 to C0307 and C0309, BioLegend). Staining was performed according to the manufacturer’s instructions. Briefly, FACS-sorted T cells were stained with 0.5 mg TotalSeq hashtag per sample for 30 min on ice and washed 5 times with PBS + 0.04% BSA. Prior to last wash, cells were passed through a 40 mm Flowmi Cell Strainer (Sigma-Aldrich) and samples were pooled. scRNAseq libraries were created according to the manufacturer’s protocol, using a 10X Chromium single cell 5’v2 chemistry kit (10X Genomics). TCRseq was library was prepared using a 10X Chromium single cell V(D)J enrichment kit for mouse T cells (10X Genomics). Nucleotide sequencing was performed on a NovaSeq6000 system (Illumina). In total, 25132 cells were analyzed for sequencing with a median sequencing-depth of 11196 reads per cell.

### Processing, integration and clustering of transcriptome data

Analysis of scRNAseq and TCRseq data was performed using CellRanger (10X Genomics, v6.0.1) and Seurat (v4.3.0) in R (v4.2.3). scRNAseq reads were aligned to the mouse reference genome mm10, and barcode counting was performed using ‘cellranger count’. TCRseq reads were aligned to the mouse reference genome mm10 and consensus TCR sequences were determined using ‘cellranger vdj’. From the gene expression count matrix, TCR and BCR genes were removed using the ‘grep’ function in R with the patterns “^Tr[abgd][cvdj]” and “Ig[hkl][cvdj]” respectively. After this, the dataset was analyzed in R using the standard Seurat pipeline. Seurat object was created using ‘CreateSeuratObject,’ with a minimum of 200 features per cell and 3 cells expressing each gene. HTO sequencing data was added as a separate assay to the Seurat object using ‘CreateAssayObject’ and HTO counts were normalized by CLR method using ‘NormalizeData.’ HTO demultiplexing was performed using ‘HTODemux’ with a positive quantile of 0.99. As a result of this, each cell in the dataset was assigned to a global (singlet, doublet, or negative) and a sample-related HTO classification. Heatmap was created of HTO demultiplexing result using ‘HTOHeatmap.’ Singlets were selected using ‘subset’, and gene expression data was normalized using ‘NormalizeData’. Variable features were determined using ‘FindVariableFeaures’ with the selection method ‘mean.var.plot’. Data was scaled using ‘ScaleData’ on variable features. Percentage of mitochondrial gene expression per cell was determined using ‘PercentageFeatureSet’ with the pattern “^mt-“. Singlets were filtered on number of RNA features (between 400 and 5000), number of RNA counts (<25000) and mitochondrial gene expression (<10%). Cell cycle scores for G2/M and S phase were assigned using ‘CellCycleScoring’, based on expression of G2/M and S phase genes. Data was normalized using ‘NormalizeData’ with normalization method ‘LogNormalize’, scale factor 25000. Top 2000 variable features were determined using ‘FindVariableFeatures’ with the selection method ‘vst’. Data was scaled using ‘ScaleData’, whereby G2M scores, S scores and percentage of mitochondrial gene expression were regressed out. Principle component analysis (PCA) was run using ‘RunPCA’ on variable features. Clustering was performed on first 20 principal components (PCs) using ‘FindNeighbors’ and ‘FindClusters’. Clustering resolution was set to 0.5. Uniform manifold approximation and projection (UMAP) visualization was performed using ‘RunUMAP’ on the first 20 PCs. Expression of known marker genes was visualized per cluster and within the UMAP projection using ‘Dotplot’, ‘VlnPlot’ and ‘FeaturePlot’ functions. Clusters with contaminating B cells (expression of *Cd19*, *Ms4a1*), APCs (expression of *Cd40*, *Cd274*, *Cd80*, *Cd86*) and low-quality cells (high percentage mitochondrial genes, low RNA counts) were removed. The pipeline was run again on filtered T cells, including determining variable features, data scaling, PCA, clustering and UMAP visualization. CD4 T-cell clusters were selected by expression of *Cd4*, in absence of expression of *Cd8a* and *Cd8b1* or TCRg/d. Within CD4 T-cell subsets, Help d5 and Help d10 were selected for further analysis. Clusters were annotated based on differentially expressed genes. Venn Diagrams were created with http://bioinformatics.psb.ugent.be/webtools/Venn/.

Previously generated scRNAseq datasets from I-A^b^-LCMV-gp66 tetramer-sorted CD4 T cells from LCMV-Armstrong or LCMV-Clone 13 derived at day 8 and day 21 post infection were obtained from (Xia et al., 2022). Feature barcode matrices derived from 10x scRNA expression libraries reads aligned to mouse genome assembly GRCm38 were imported into R (version 4.4.0). A Seurat-based (version 5.2.1) quality control on the scRNA-seq samples was performed individually. TCR and BCR counts were removed from the raw gene expression count matrix (UMI) to prevent TCR/BCR chain driven clustering bias. A per-sample Seurat object was created by selecting cells on the criterium of containing a minimum of 200 genes and genes on being expressed in a minimum of 3 cells. Cells (∼30%) were excluded from the subsequent analysis based on the mitochondrial content (>5%), RNA count (<1000 and >25000) and genetic content (<400 and >4000) The top 2000 highly variable genes were selected using ‘FindVariableFeatures’ for subsequent analysis. Cells of all 19 samples were normalized and scaled individually while regressing out mitochondrial gene expression and cell cycle scores for G2/M and S phase genes assigned using ‘CellCycleScoring’ as described above. Clustering was done on the first 30 principal components (PCs) with the default cluster resolution. Doublets were removed using ‘DoubletFinder’ based on the first 40 principal components (PCs).

Singlets of each sample were integrated by identifying anchors between datasets, using the top 2000 variable genes, 30 reciprocal PCA dimensions and an integration strength of 5. Clusters with contaminating B cells (expression of Cd19 Ms4a1), CD8 T cells (expression of Cd8a, Cd8b), APCs (expression of Cd40, Cd80, Cd86) or low-quality cells were removed. Data scaling and PCA was done once more on the integrated and filtered object. A k-nearest neighbors graph was generated with the first 20 PCs and community detection for cluster identification was performed using the Louvain algorithm with a resolution of 0.4. The integrated matrix visualized as an UMAP was subsequently divided based on the day of infection (Day 8 or Day 21) and the viral strain (LCMV-Arm and LCMV-c13) as annotated in the metadata of the Seurat object.

Clusters were annotated based on differentially expressed genes. Subsequently, clusters were compared to CD4 T cell-specific signatures derived from Th1, Tfh or Th1/Tfh precursors, all derived from vaccine induced CD4 T cells from this article, or to a curated stem-like signature (Cardenas et al., 2024) to enable cluster annotation via scoring based on gene signature module scores, which were calculated using Seurat’s ’AddModuleScore’ function by averaging the expression of the signature genes for each cell. cluster defining markers were selected and the relative normalized expression values represented in z-scores were visualized in a heatmap using Seurat’s ‘pheatmap’.

### Statistics

For mouse experiments, statistical analysis was performed with GraphPad Prism 9.3.1 (Dotmatics) using two-tailed unpaired Student *t-*test when comparing two groups, One-Way ANOVA with Dunnett post hoc test to correct for multiple testing when comparing multiple groups at one timepoint or Two-Way ANOVA with Šidák post hoc test to correct for multiple testing when comparing multiple groups at multiple time points. P values < 0.05 were considered statistically significant. Statistics were denoted as follows; *; P≤0.05, **; P≤0.01, ***; P≤0.001 and ****; P≤0.0001. Plots were generated with ggplot2 (3.3.3) and GraphPad Prism 9.3.1.

## Author contributions

Conceptualization: D.M.T.B, J.Bo, F.S. Methodology and investigation: D.M.T.B, J.Bu, M.D.S, X.L, T.d.W, Y.X, F.S. Formal analysis, visualization and data curation: D.M.T.B, J.Bu, M.d.K, F.S. Writing - original draft: D.M.T.B, J.Bo, F.S. Writing - review and editing: D.M.T.B, J.Bu, M.D.S, M.d.K, X.L, T.d.W, Y.X, J.Bo, F.S. Project administration, funding acquisition and supervision: J.Bo. and F.S.

## Supporting information

Supplemental document contains 7 supplementary figures

## Acknowledgements

We thank the Animal facility, Flow Cytometry Facility and Leiden Genome Technology Center from the LUMC for technical assistance. This work was supported by grants 11079 (to J.B. and Y.X) and 10894 (to J.B.) from the Dutch Cancer Society (KWF); a grant from Oncode to JB, and a VENI grant (ZonMw project n. 09150162010046) from the Dutch Science Foundation (NWO) to F.S.

## Declaration of interests

The authors declare no competing interests.

## Supplemental information

- Document S1: Supplementary figures 1-7.

## Notes

### Competing Interest Statement

The authors have declared no competing interest.

### Summary of Updates

The addendum reinstates omitted text in the sections: - Specific costimulatory signals driving formation of the common Th1/Tfh precursor pool and subsequent Th1- or Tfh differentiation - Th1/Tfh precursors differentiate into either Th1 or Tfh cells in a second priming step on cDC1s or B cells, respectively.

