## Supplemental document contains 7 supplementary figures for "Requirements for development of T-helper1 and T-follicular helper cells from a common precursor"

*Shared first authors

**Shared last authors

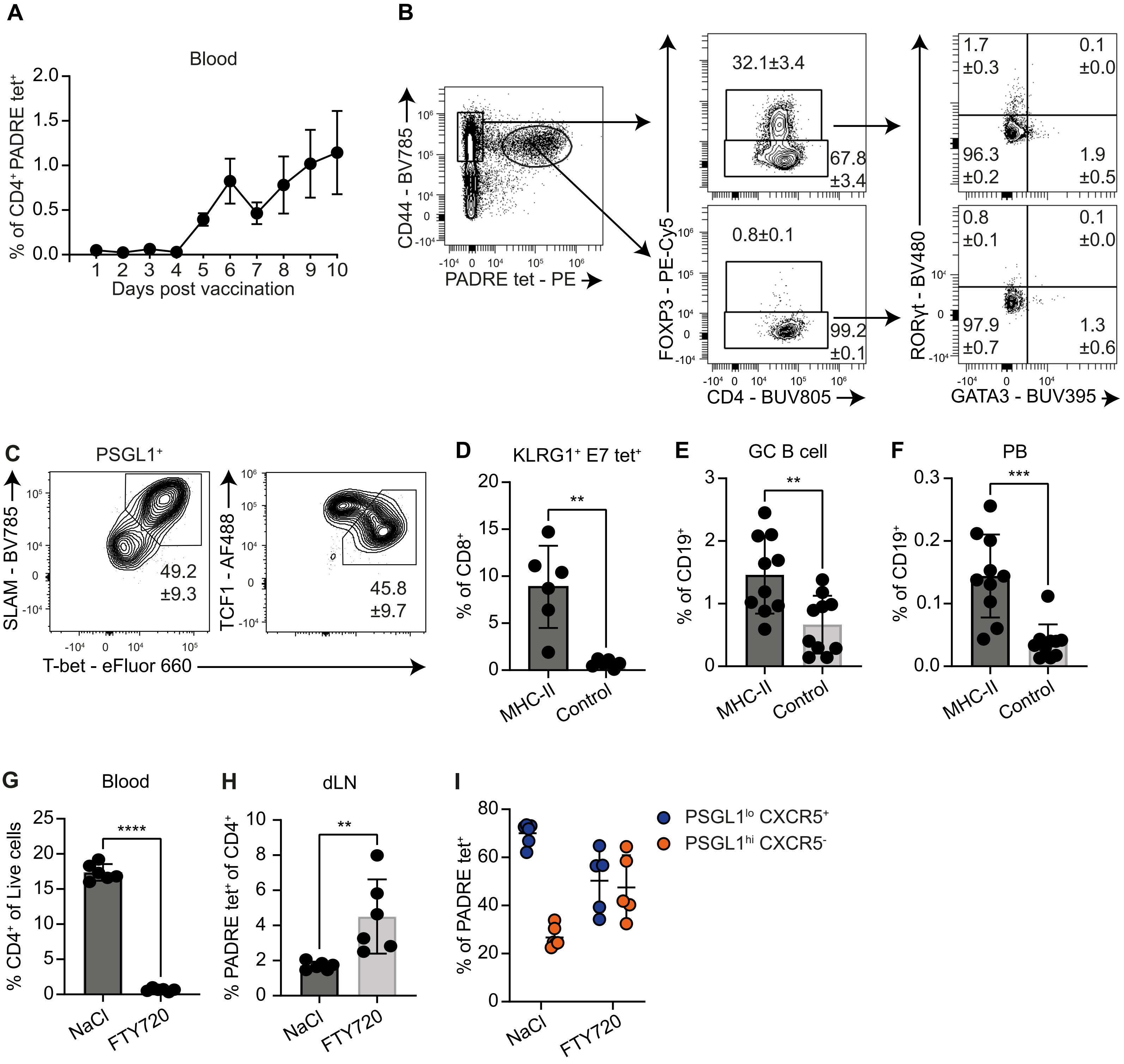


**Supplementary Figure 1: A vaccination model to study differentiation of vaccine-primed CD4 T cells into Th1 and Tfh cells.** **A)** Frequency of PADRE tet$^+$ CD4 T cells in blood over time after vaccination (N=5). **B)** Representative contour plots depicting Foxp3, GATA3 and RORγt expression in CD44$^+$ PADRE tet$^-$ (top) and CD44$^+$ PADRE tet$^+$ (bottom) CD4 T cells. **C)** SLAM, T-bet and TCF1 expression by PSGL1$^+$ CXCR5$^-$ PADRE tet$^+$ CD4 T cells. **D)** Frequency of antigen-specific effector CD8 T cells (E7 tet$^+$ KLRG1$^+$) in blood at day 10 after MHC-II or control vaccination. **E)** Frequency of GL7$^+$ FAS$^+$ GC B cells in the dLN at day 8 post-vaccination (N=10 per group). Cells were pre-gated as CD19$^+$ IgD$^-$ live cells. **F)** Frequency of plasmablasts (PB; CD19$^{int/lo}$ IgD$^-$ CD138$^+$ TACI$^+$) at day 8 days after vaccination in the dLN (N=10 mice per group). **G-I)** Mice were treated with FTY720 or vehicle at day 0, 3 and 6 after vaccination and analyzed at day 7 (N=6 per group). **G)** Frequency of total CD4 T cells in blood. **H)** PADRE tet$^+$ CD4 T cells in dLN. **I)** Frequency PSGL1$^{lo}$ CXCR5$^+$ and PSGL1$^{hi}$ CXCR5$^-$ PADRE tet$^+$ CD4 T cells in dLN. All graphs depict mean ± SD; D-H) Unpaired student-t test.

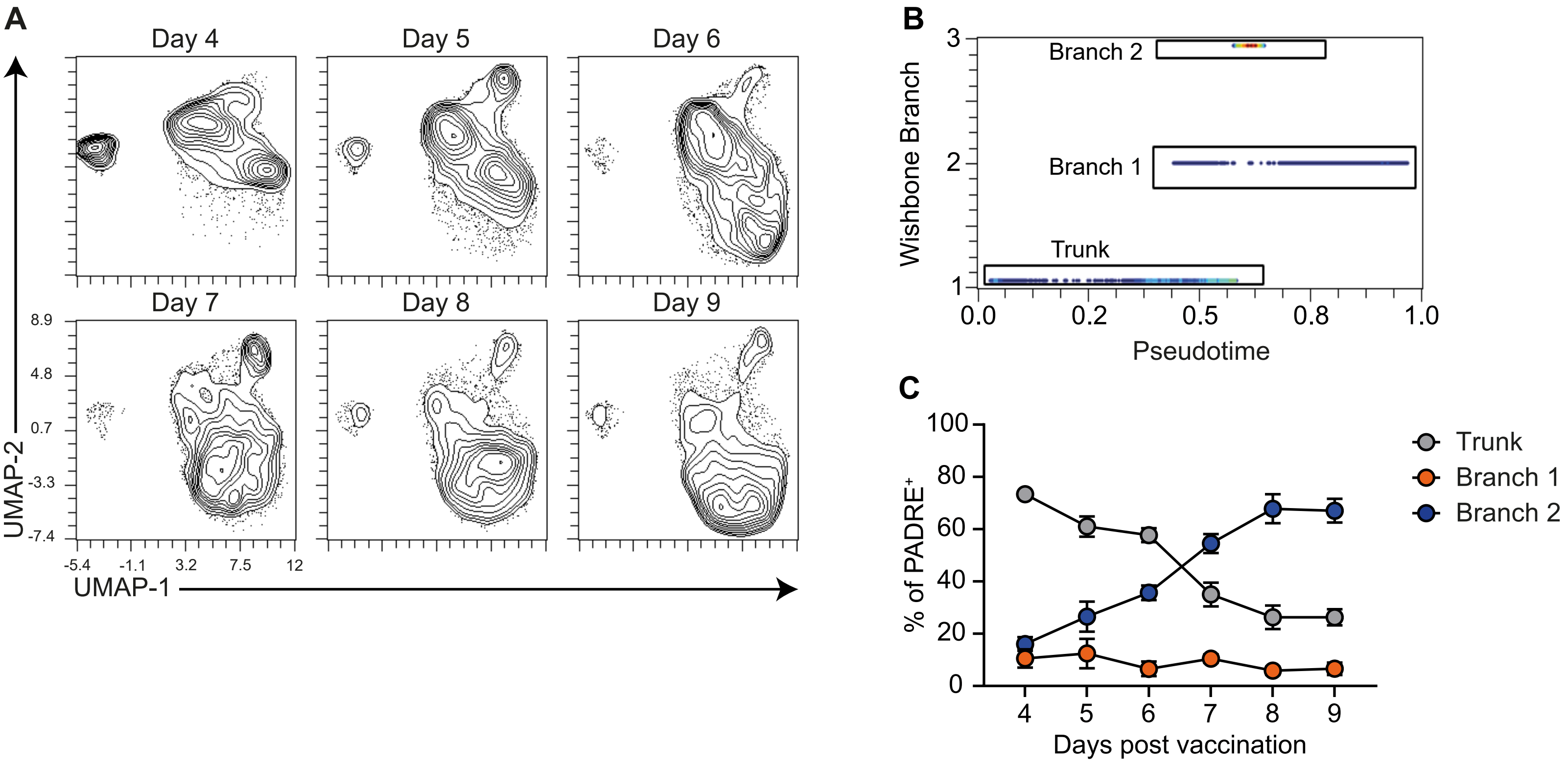


**Supplementary Figure 2: Temporal analysis highlights a common differentiation branchpoint for Th1 and Tfh** **A)** UMAP projections showing PADRE tet$^+$ CD4 T cells in the dLN from day 4-9 post-vaccination (N=5 per time point). **B)** Wishbone trajectory and branching of concatenated PADRE tet$^+$ CD4 T cells. **C)** Frequency of PADRE tet$^+$ CD4 T cells within trunk, branch 1 and branch 2 in the dLN as calculated by Wishbone. All graphs depict mean ± SD.

A

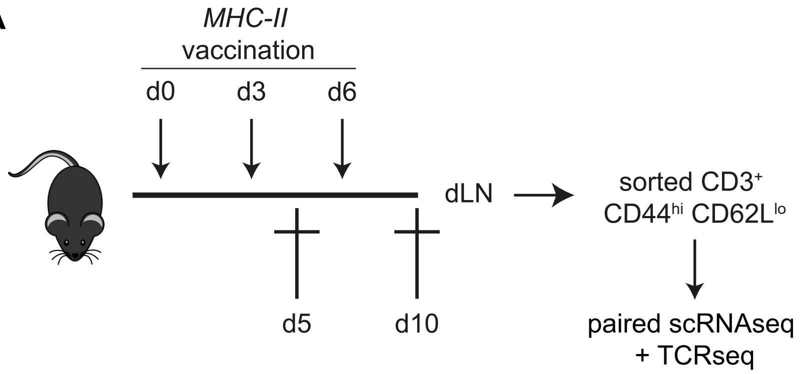


B

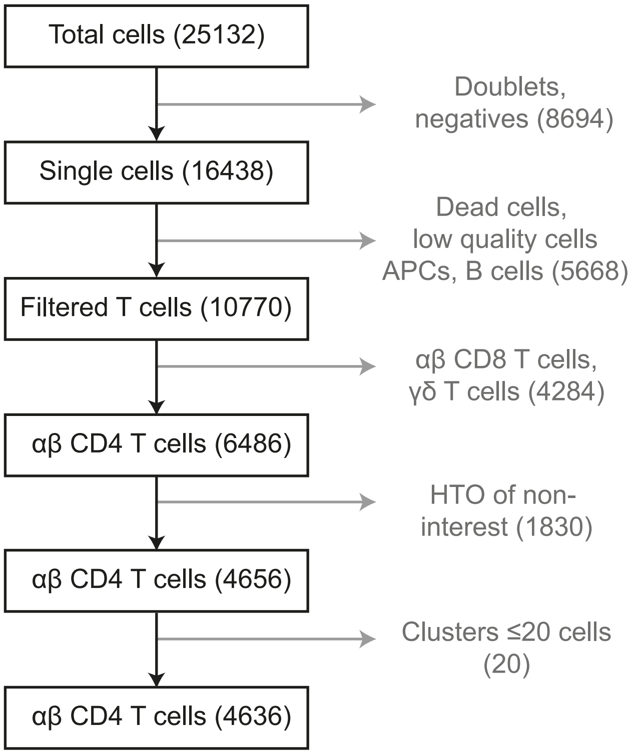


C

| | Cell number per cluster | | |
|---|---|---|---|
| | *d5* | *d10* | *total* |
| *cl.0* | 721 | 412 | 1133 |
| *cl.1* | 182 | 881 | 1063 |
| *cl.2* | 258 | 413 | 671 |
| *cl.3* | 315 | 299 | 614 |
| *cl.4* | 254 | 259 | 513 |
| *cl.5* | 310 | 156 | 466 |
| *cl.6* | 176 | 35 | 176 |

D

| | Expanded TCR clones per cluster | | |
|---|---|---|---|
| | *unique* | *shared* | *total* |
| *cl.0* | 45 | 37 | 82 |
| *cl.1* | 99 | 117 | 216 |
| *cl.2* | 22 | 95 | 117 |
| *cl.3* | 0 | 3 | 3 |
| *cl.4* | 0 | 20 | 20 |
| *cl.5* | 15 | 85 | 100 |
| *cl.6* | 0 | 13 | 13 |

E

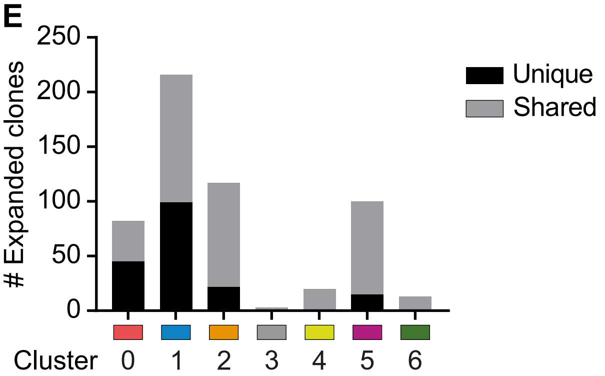


**Supplementary Figure 3: scRNAseq and TCRseq of dLN-derived CD4 T cells after vaccination A-B)** Schematic overview of mouse treatment, sample preparation (A) and pipeline for quality control and filtering steps of scRNAseq data analysis (B). **C, D)** Tables report numbers of cells identified per cluster (cl.) (C) at sample days 5 and 10, or numbers of expanded TCR clones per cluster, which identify clones that were uniquely found in one cluster, or were shared between at least two clusters (D). **E)** Number of expanded TCR clones (>1 cell per clone) per cluster. Black: TCR clones unique to one cluster; grey: TCR clones shared between two or more clusters.

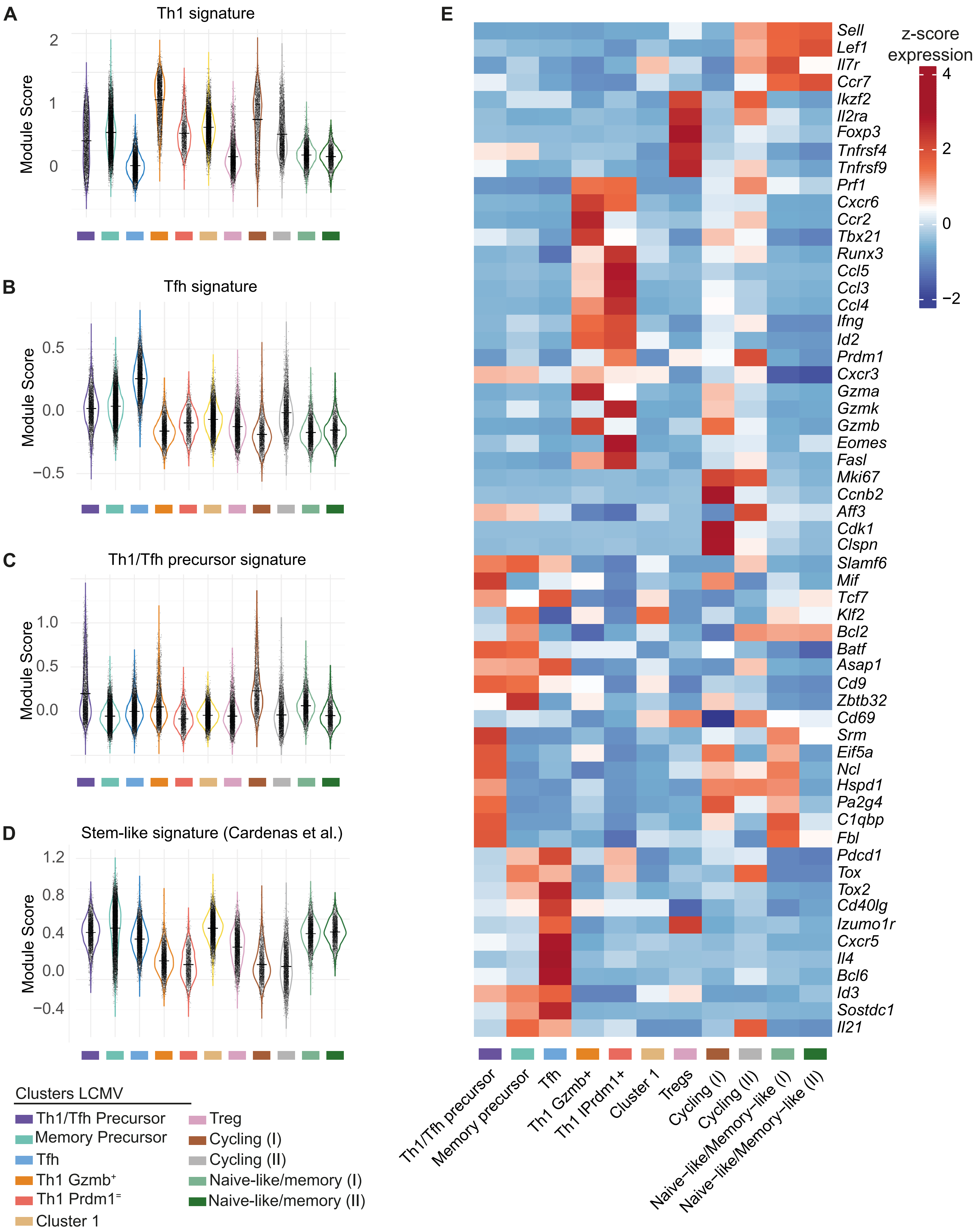
A
Th1 signature
Module Score
B
Tfh signature
C
Th1/Tfh precursor signature
D
Stem-like signature (Cardenas et al.)
E
z-score expression
Sell
Lef1
Il7r
Ccr7
Ikzf2
Il2ra
Foxp3
Tnfrsf4
Tnfrsf9
Prf1
Cxcr6
Ccr2
Tbx21
Runx3
Ccl5
Ccl3
Ccl4
Ifng
Id2
Prdm1
Cxcr3
Gzma
Gzmk
Gzmb
Eomes
Fasl
Mki67
Ccnb2
Aff3
Cdk1
Clspn
Slamf6
Mif
Tcf7
Klf2
Bcl2
Batf
Asap1
Cd9
Zbtb32
Cd69
Srm
Eif5a
Ncl
Hspd1
Pa2g4
C1qbp
Fbl
Pdcd1
Tox
Tox2
Cd40lg
Izumo1r
Cxcr5
Il4
Bcl6
Id3
Sostdc1
Il21
Th1/Tfh precursor
Memory precursor
Tfh
Th1 Gzmb+
Th1 IPrdm1+
Cluster 1
Tregs
Cycling (I)
Cycling (II)
Naive-like/Memory-like (I)
Naive-like/Memory-like (II)
Clusters LCMV
Th1/Tfh Precursor
Memory Precursor
Tfh
Th1 Gzmb⁺
Th1 Prdm1⁼
Cluster 1
Treg
Cycling (I)
Cycling (II)
Naive-like/memory (I)
Naive-like/memory (II)

**Supplementary Figure 4: Gene profile of the LCMV-specific CD4 T cells**

**A-C)** Violin plots depicting mean for module score of expression levels of Th1 (A), Tfh (B) and Th1/Tfh precursor (C) signatures derived from vaccination in total GP-66 tet$^{+}$ LCMV-specific CD4 T cells derived from LCMV-Armstrong and Clone13 at day 8 and day 21 derived from (Xia et al., 2022) per cell cluster. **D)** Violin plots depicting mean for module score of expression levels of curated Top genes for stem-like cells in LCMV-specific CD4 T cells (Cardenas et al., 2024) in total GP-66 tet$^{+}$ LCMV-specific CD4 T cells derived from LCMV-Armstrong and Clone13 at d8 and d21 per cell cluster. E) Heatmap of top differentially expressed genes for LCMV-specific CD4 T cells. Colors represent normalized (z-scored) expression per gene among each cluster.

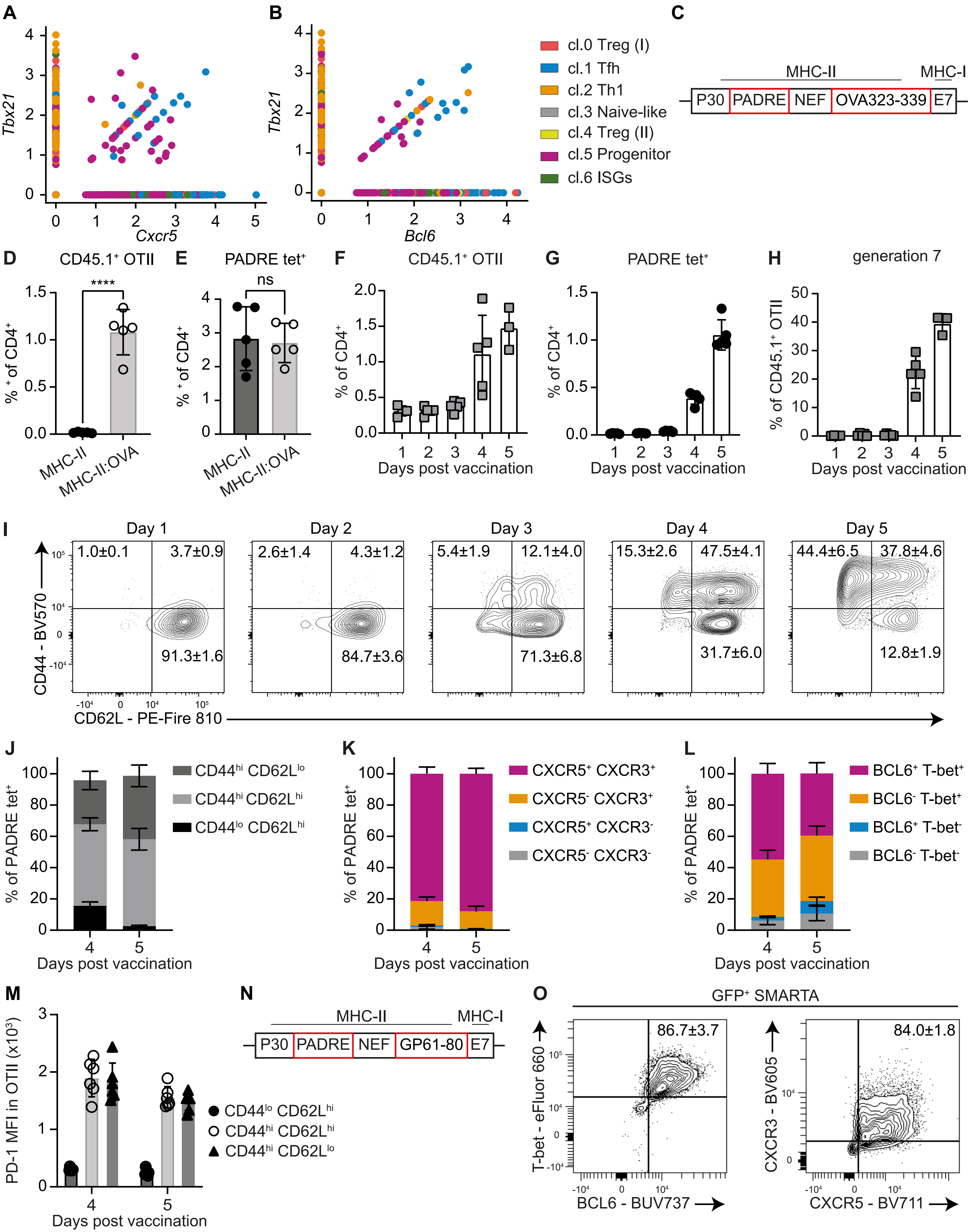
A
B
C
Tbx21
Cxcr5
Bcl6
cl.0 Treg (I)
cl.1 Tfh
cl.2 Th1
cl.3 Naive-like
cl.4 Treg (II)
cl.5 Progenitor
cl.6 ISGs
MHC-II
MHC-I
P30
PADRE
NEF
OVA323-339
E7
D
CD45.1⁺ OTII
****
% ⁺ of CD4⁺
MHC-II
MHC-II:OVA
E
PADRE tet⁺
ns
F
CD45.1⁺ OTII
% of CD4⁺
Days post vaccination
G
PADRE tet⁺
H
generation 7
% of CD45.1⁺ OTII
I
Day 1
Day 2
Day 3
Day 4
Day 5
1.0±0.1
3.7±0.9
91.3±1.6
2.6±1.4
4.3±1.2
84.7±3.6
5.4±1.9
12.1±4.0
71.3±6.8
15.3±2.6
47.5±4.1
31.7±6.0
44.4±6.5
37.8±4.6
12.8±1.9
CD44 - BV570
CD62L - PE-Fire 810
J
% of PADRE tet⁺
CD44hi CD62Llo
CD44hi CD62Lhi
CD44lo CD62Lhi
K
CXCR5⁺ CXCR3⁺
CXCR5⁻ CXCR3⁺
CXCR5⁺ CXCR3⁻
CXCR5⁻ CXCR3⁻
L
BCL6⁺ T-bet⁺
BCL6⁻ T-bet⁺
BCL6⁺ T-bet⁻
BCL6⁻ T-bet⁻
M
PD-1 MFI in OTII (x10³)
CD44lo CD62Lhi
CD44hi CD62Lhi
CD44hi CD62Llo
N
MHC-II
MHC-I
P30
PADRE
NEF
GP61-80
E7
O
GFP⁺ SMARTA
86.7±3.7
84.0±1.8
T-bet - eFluor 660
BCL6 - BUV737
CXCR3 - BV605
CXCR5 - BV711

**Supplementary Figure 5: Phenotypic analysis of the common Th1/Tfh precursor pool.**

**A-B)** Expression levels per cell of Tbx21 and Cxcr5 (A) or Tbx21 and Bcl6 (B) mRNA. CD4 T cell clusters from scRNAseq are color-coded as in Figure 2. **C)** Schematic overview of epitopes in MHC-II:OVA vaccine. **D, E)** Frequency of CD45.1$^{+}$ OTII (D) or PADRE tet$^{+}$ (E) CD4 T cells in the dLN of mice at day 8 after vaccination with MHC-II or MHC-II:OVA plasmid (N=5 per group), as determined by flow cytometry. Unpaired student's-t test. **F, G)** Frequency of CD45.1$^{+}$ OTII (F) or CD45.2$^{+}$ PADRE tet$^{+}$ (G) CD4 T cells at indicated times after vaccination, as determined by flow cytometry. **H)** Frequency of CD45.1$^{+}$ OTII that underwent 7 cell divisions at indicated time points post-vaccination (N=3-5 per timepoint), as determined by CFSE dilution. **I**) Flow cytometric detection of CD44 and CD62L on CD45.1+ OTII cells in the dLN at indicated time points. **J-L)** Graphs displaying mean ± SD of CD44 and CD62L (K), CXCR5 and CXCR3 (L), or BCL6 and T-bet (M) protein levels as detected by flow cytometry in PADRE tet$^{+}$ CD4 T cells at days 4 and 5 post-vaccination (N=5 per time point). **M)** Median fluorescent intensity (MFI) for PD-1 on CD44$^{lo}$ CD62L$^{hi}$ CD44$^{hi}$, CD62L$^{hi}$ and CD44$^{hi}$ CD62L$^{lo}$ CD45.1$^{+}$ OTII cells at day 4 or d5 after vaccination. **N)** Schematic overview of epitopes in the LCMV-GP$_{61-80}$ modified MHC-II vaccine. **O)** Representative contour plots depicting T-bet and BCL6 or CXCR3 and CXCR5 protein levels in GFP$^{+}$ SMARTA cells at day 4 post-vaccination (N=6 per group). All graphs depict mean ± SD.

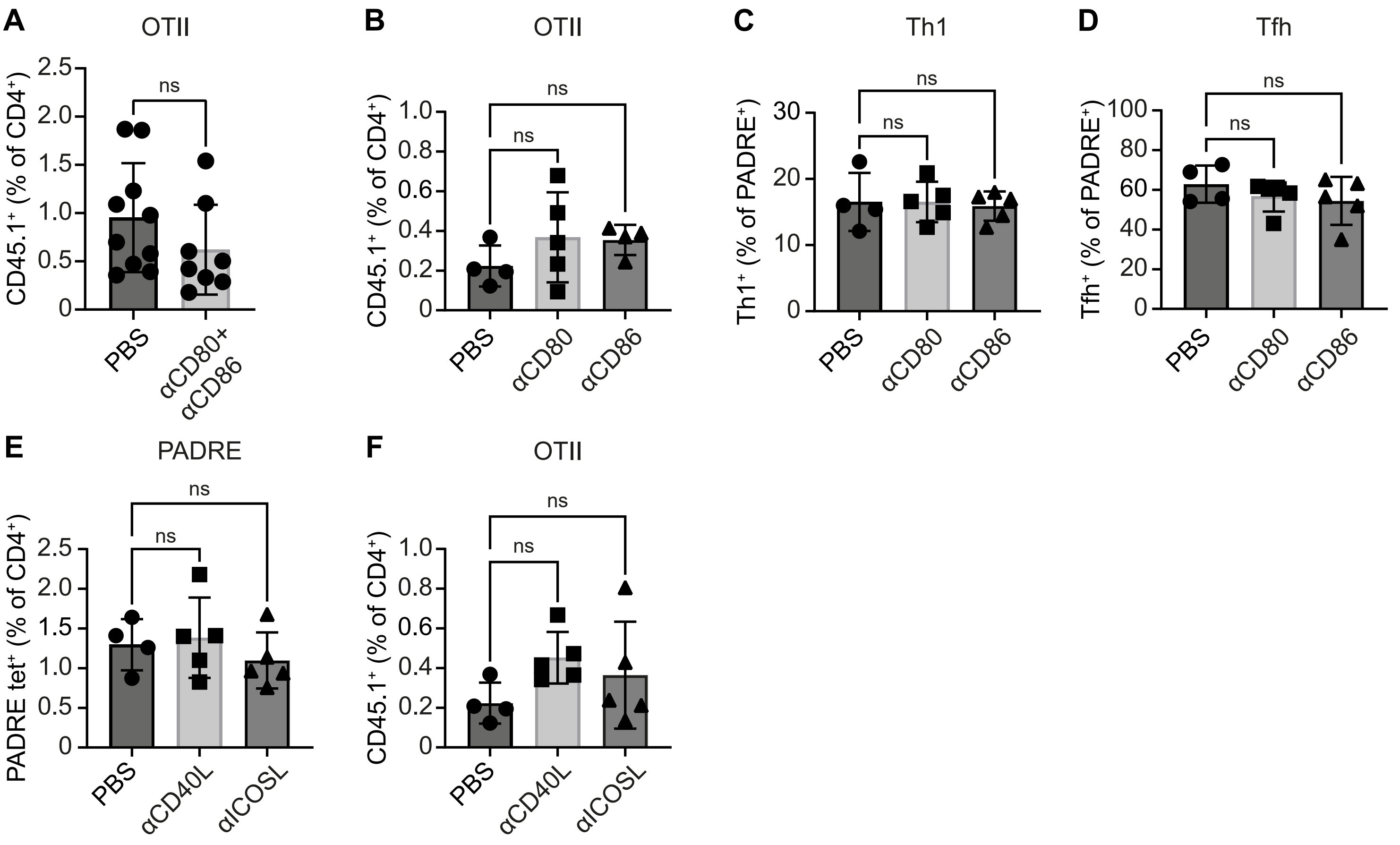


**Supplementary Figure 6: Costimulatory molecules driving CD4 T-cell differentiation.**
Mice received OTII cells and MHC-II:OVA vaccine and were treated with blocking mAbs to the indicated molecules or PBS (control). At day 5 after vaccination, dLNs were harvested and cells were analyzed by flow cytometry. **A)** Quantification of $CD45.1^{+}$ OTII cells in dLNs after blocking CD80 and CD86 or control (N=9-10 per group pooled from two experiments). Unpaired student's-t test. **B-D)** Quantification of $CD45.1^{+}$ OTII cells (B), $SLAM^{+}$ $Tbet^{+}$ Th1 PADRE $tet^{+}$ cells (C) and $CXCR5^{+}$ $PD\text{-}1^{+}$ Tfh PADRE $tet^{+}$ cells (D) in dLN after blocking either CD80, or CD86, or control. **E-F)** Quantification of PADRE $tet^{+}$ (E) and CD45.1 OT-II (F) cells in dLN after blocking CD40L or ICOSL, or control (N=4-5 per group). B-F) One-Way ANOVA with Dunnett post hoc test was used to correct for multiple testing when comparing inhibitory conditions to control. Data are representative of at least two independent experiments. **A-F)**. All graphs depict mean ± SD.

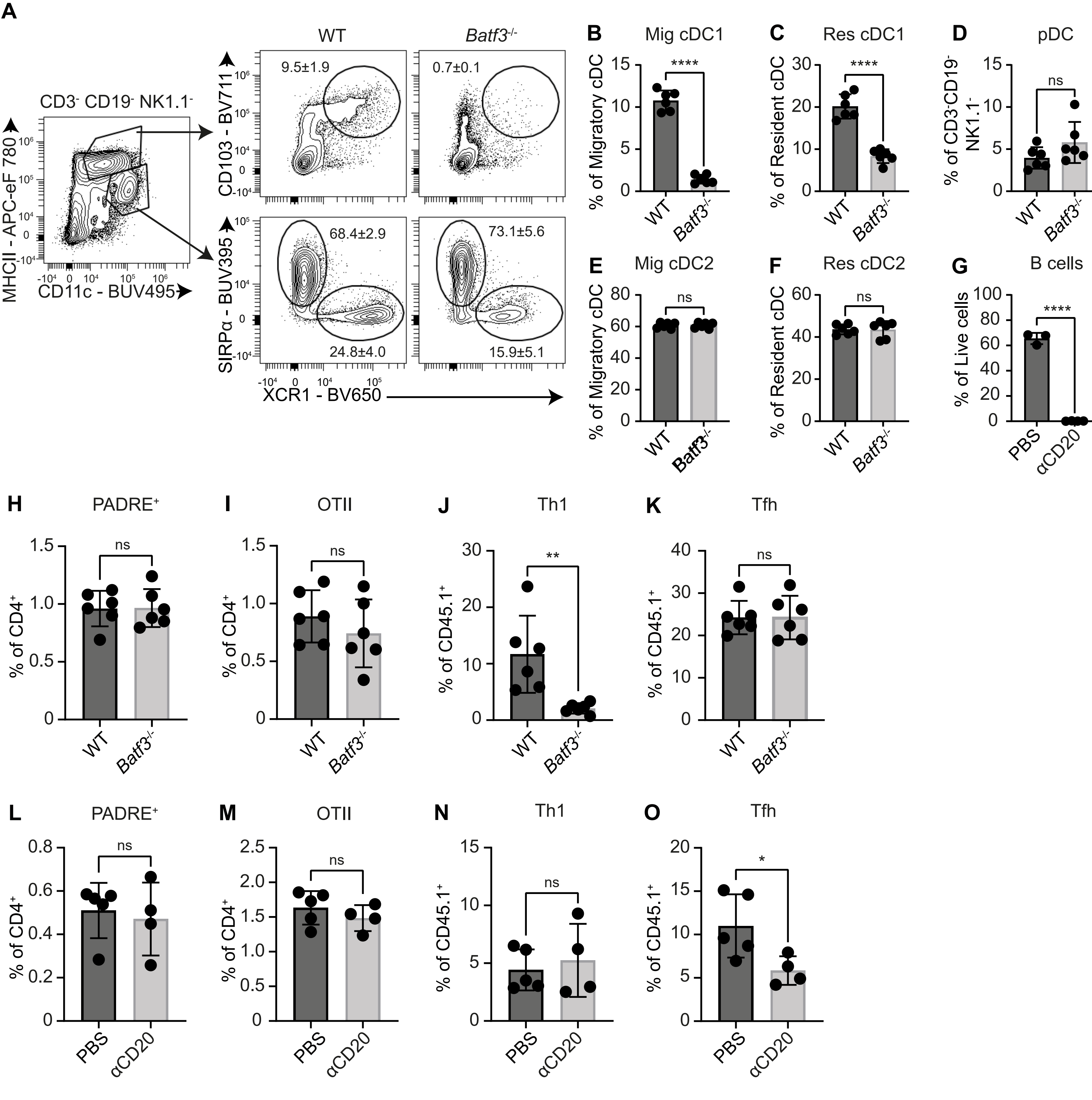


**Supplementary Figure 7: Antigen presenting cells driving CD4 T-cell differentiation.**
WT or *Batf3*$^{-/-}$ mice received OTII cells and MHC-II:OVA vaccine and were treated or not with depleting anti-CD20 mAb or PBS (control) as indicated. At day 5 after vaccination, dLNs were harvested and cells were analyzed by flow cytometry. **A)** Gating strategy to identify migratory cDC1s (MHCII$^{hi}$ CD11c$^{int}$ CD103$^{+}$ XCR1$^{+}$), resident cDC1s (MHCII$^{int}$ CD11c$^{hi}$ XCR1$^{+}$ SIRPα$^{-}$) and resident cDC2s (MHCII$^{int}$ CD11c$^{hi}$ XCR1$^{-}$ SIRPα$^{+}$) in the dLN at day 3 post-vaccination. **B-F)** Quantification of migratory (B) and resident (C) cDC1, pDCs (Ly6C$^{+}$ Siglec H$^{+}$; D), migratory (MHCII$^{hi}$ CD11c$^{int}$ XCR1$^{-}$ SIRPα$^{+}$; E) and resident cDC2 (F) in WT and *Batf3*$^{-/-}$ mice (N=6 per group). **G)** Quantification of CD19$^{+}$ B cells in peripheral blood two days after treatment with anti-CD20 mAb (N=3-4 per group). **H, L)** Frequency of PADRE tet$^{+}$ among CD4 T cells in dLNs of indicated test groups. **I, M)** Frequency of CD45.1$^{+}$ OTII cells in dLN of indicated test groups. **J-K, N-O)** Frequency of CD45.1$^{+}$ OTII cells with SLAM$^{+}$ T-bet$^{+}$ Th1 phenotype (J, N) or CXCR5$^{+}$ PD-1$^{+}$ Tfh phenotype (K, O) CD45.1$^{+}$ OTII cells in dLNs of indicated test groups. All graphs depict mean ± SD (N=6 per group in A-K; N=4-5 per group in L-O). Unpaired student's-t test.
